# Automated Discovery of Therapeutic Biomaterial for Renally Impaired Hyperuricemia Patients by Natural Language Processing and Machine Learning

**DOI:** 10.1101/2025.03.05.641578

**Authors:** Xiaodong Zeng, Jiahao Qiu, Xin Zhao, Kangfei Liu, Longfei Zhao, Junkang An, Liang Xiang, Tianzhi Liu, Zhichao Wang, Weidi Xie, Mengdi Wang, Jie Luo, Shiyi Zhang

## Abstract

The exponential growth of scientific publications presents opportunities for researchers to identify valuable knowledge, especially in the highly interdisciplinary field --- biomaterials, where exploiting possible connections between unmet clinical needs and materials properties from literatures is crucial. However, with traditional literature reading, it is extremely challenging to marry unmet clinical needs with existing materials reported for different applications or other purposes. Here, to provide a not-renally cleared therapeutics for renally impaired hyperuricemia patients, we designed a multi-tiered framework MatWISE that fuses state-of-the-art natural language processing, semantic relationship mapping, and machine learning to automate the complex process of material discovery from a sea of scientific literatures published until December of 2022, and successfully identified and optimized δ-MnO_2_ into an orally administered, nonabsorbable uric acid (UA) lowering biomaterial. δ-MnO_2_ had superior serum and urine UA-lowering effect in three hyperuricemia mouse models, by comparing with a standard of care drug. δ-MnO_2_ is highly promising to serve as a safe and effective UA-lowering drug for renally impaired hyperuricemia patients. We demonstrated a new research paradigm for biomaterials that combining state-of-the-art machine learning techniques and a handful of experiments to discover a translationally relevant material from the massive existing research, for an unmet clinical need.

## 1. Introduction

In recent years, when the scientific literature has grown exponentially, a plethora of literature has provided opportunities but also difficulties for interdisciplinary researchers to comprehend and utilize literature knowledge.^[1]^ Human cognitive limitations often lead to biased selections of personally familiar materials, for various applications, thereby preventing fair selection and thorough comparison among all possible solutions. In parallel, the explosive number of materials reported in literatures have not been fully appreciated for the translational medicine purpose. Consequently, there is a pressing need for an innovative research paradigm for biomaterials. Recent progress in artificial intelligence (AI),^[2–7]^ offers an unprecedented opportunity to mine latent, clinically relevant information from the wealth of existing literature, potentially bridge the gap of material properties and unmet clinical need’s requirements.

With the revolutionary changes in the NLP field brought about by Generative Pretrained Transformer (GPT) type language models and Large Language Models (LLMs),^[8]^ there exists a tantalizing prospect of directly sourcing answers through automated language generation. However, while these models excel in language generation, they are often constrained by the pitfalls of hallucination and limited parameter spaces, rendering them less suitable for precise material screening.^[9]^ Traditional NLP techniques such as word2vec^[7]^ has been attempted used to capture the hidden relationship between domain specific entities. However, they often yield irrelevant co-occurring entities (Figure S1a, Supporting Information) and miss logically related information that’s not in close textual proximity (Figure S1b, Supporting Information). BioGPT,^[10]^ a specialized language model explicitly trained on biomedical literature, embeddings from which could capture long range associations offering the potential to facilitate the resolution of the above-mentioned challenges, and ultimately enable the discovery of novel biomaterials. To further refine the screening process, we integrate GPT-4 with web search augmentation and domain-specific databases. This enhancement allows for the extraction of well-documented properties of material candidates, thereby streamlining the subsequent experimental verification phase. To our knowledge, this is the first comprehensive workflow— termed MatWISE (Materials Workflow for Intelligent Screening and Evaluation)—that leverages state-of-the-art NLP and machine learning technologies for the intelligent screening and evaluation of materials aimed at addressing unmet clinical needs within the expansive landscape of existing literature.

Hyperuricemia is a risk factor for gout.^[11,12]^ Moreover, hyperuricemia also contributes to the progression of chronic kidney disease (CKD), obesity, hypertension, type 2 diabetes and coronary artery disease.^[12–14]^ There are three therapeutic strategies by lowering UA production or increasing UA elimination, including xanthine oxidase inhibitors (XOIs, allopurinol, topiroxostat, and febuxostat), uricosuric agents (probenecid, lesinurad, and benzbromarone), exogenous UA oxidase (rasburicase and pegloticase) (Figure 1a).^[15]^ However, these medications are not suited for renally impaired patients, like CKD patients, as their systemic exposure and renal clearance would exacerbate renal burden.^[16]^

**Figure 1.**
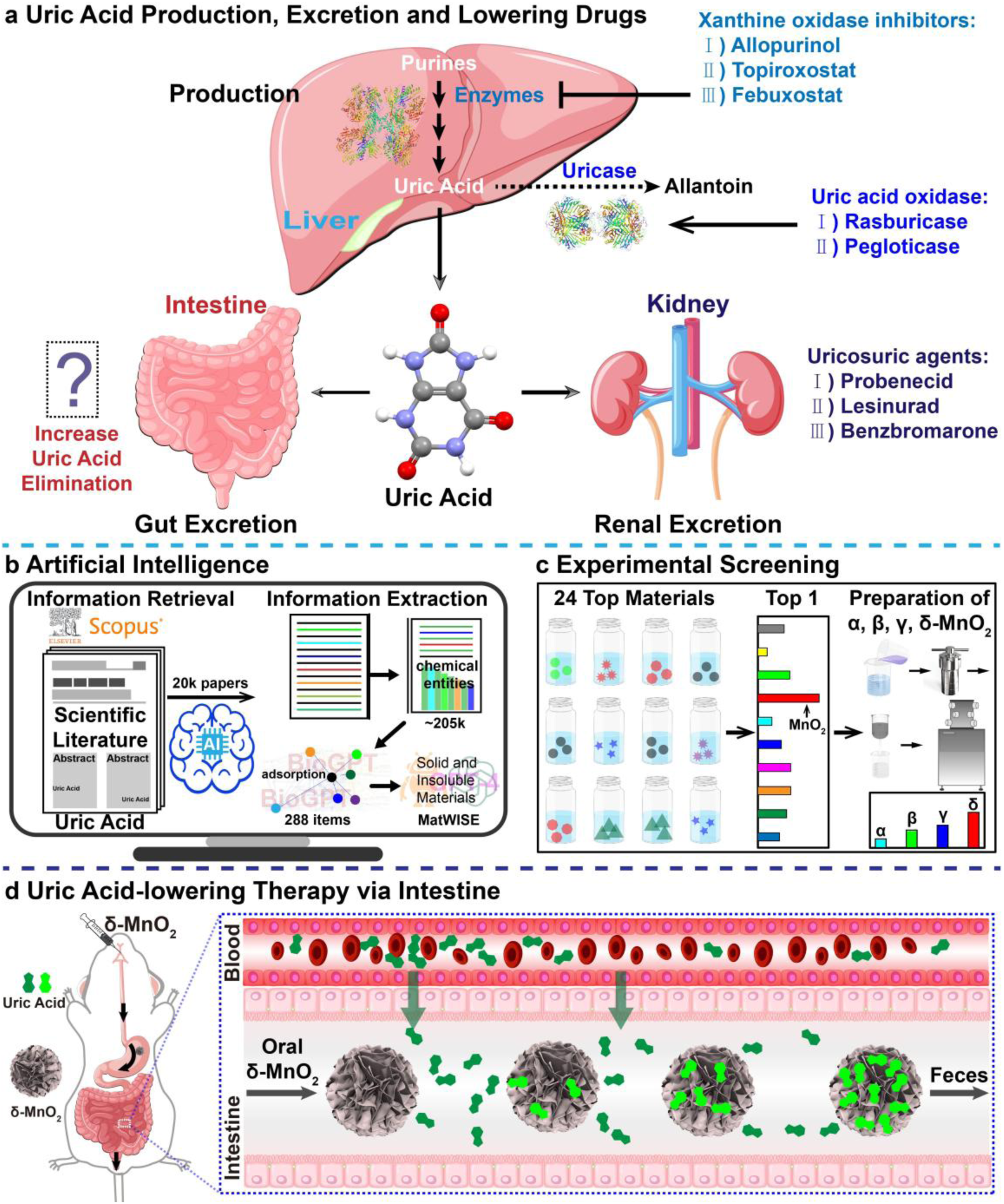
UA metabolism, approved lowering drugs and the schematic illustration of NLP-assisted discovery of nonabsorbable δ-MnO_2_ for hyperuricemia therapy via removing UA from intestine. a) The scheme for UA production, excretion pathway and approved UA-lowering drugs. b) MatWISE-Material Workflow for Intelligent Screening and Evaluation. c) Experiment validation for the UA removal ability of materials. d) The scheme of oral δ-MnO_2_ for UA-lowering therapy via UA removal from intestine.

90% of hyperuricemia are caused by insufficient UA excretion.^[17]^ Around two-thirds of UA is excreted by the kidneys, and the remaining one-third (about 200 mg per day) is eliminated by the gut.^[18]^ Furthermore, the fraction of intestinal UA removal may have a compensatory increase to 50-70% in CKD patients due to the decline of renal clearance.^[19]^ Oral administration of sevelamer, activated charcoal, montmorillonite, uricase, and bacteria have been attempted for UA adsorption or degradation in the gastrointestinal (GI) tract to reduce blood UA levels, while low efficiency, easy deactivation, and potential microbiota dysbiosis limit their clinical translation.^[19–26]^ Therefore, the development of a safe and highly efficient material to adsorb or degrade UA through GI tract might aid in meeting the clinical need for renally impaired hyperuricemia patients.

Herein, we discovered that manganese dioxide (MnO_2_) has the capability to remove UA efficiently after tapping the latent materials with semantic relationships of UA adsorption embedded within extant research corpora in massive UA related literatures assisted by MatWISE (Material Workflow for Intelligent Screening and Evaluation) (Figure 1b). Studied on four kinds of MnO_2_, showing that the δ-MnO_2_ had the highest UA removal capacity through adsorption and oxidation (Figure 1c). There are negligible manganese intake and no observable adverse effects in normal mice after repeated oral administration of δ-MnO_2_. Finally, we evaluated the UA-lowering capacity of δ-MnO_2_ in three hyperuricemia mouse models, revealing the outstanding effect of decreasing serum UA of δ-MnO_2_ and an improved ability to diminish urinary UA than allopurinol. It could provide a solution of unmet clinical needs for CKD patients. Additionally, it has the potential to reduce drug dosage and associated clinical risks when combined with other approved UA-lowering medications. Our findings reveal that oral administration of MnO_2_ effectively lowers hyperuricemia in mice by adsorbing and degrading UA in the GI tract (Figure 1d) and verify the feasibility of developing new drugs assisted by artificial intelligence.

## 2. Results and Discussion

### 2.1. Data-driven discovery of UA adsorption materials

We developed a pipeline termed MatWISE, to automate the search and selection process. We first established a database of ∼4.5k chemical entities related to UA using the state-of-the-art NLP techniques from 20k literatures (Figure 2a). In detail, literatures presenting the phrase “uric acid” anywhere in their titles and abstracts were retrieved and extracted from Scopus. To thoroughly identify all possible materials, we used a pretrained neural network model Stanza Python NLP library for biomedical and clinical text, with the aid of BioNLP13CG corpus, to extract 16 types of ∼524k scientific entities with ∼205k chemical terms (Figure 2b).^[27]^ Subsequently, individual chemical entities were connected based on entity linking (EL) to their corresponding International Union of Pure and Applied Chemistry (IUPAC) name to ensure proper disambiguation and create a chemical entities database of ∼4.5k entries. To extract latent associations between entities, biomedical NLP models (e.g. BioBERT (2019),^[28]^ SciBERT (2019),^[29]^ PubMedBERT (2021),^[30]^ and BioGPT (2022)^[10]^), could be used. Here we leveraged BioGPT, a most recently developed domain-specific generative transformer language model pre-trained on a vast biomedical literature of 15M PubMed abstracts, not for its language generation capability to answer domain-specific questions —which has limitations such as hallucinations,^[9]^ limited parameter space, and difficulty in comprehending advanced questions^[8]^— but for its ability to provide valuable embeddings that capture semantic relationships in high-dimensional vector spaces.

**Figure 2.**
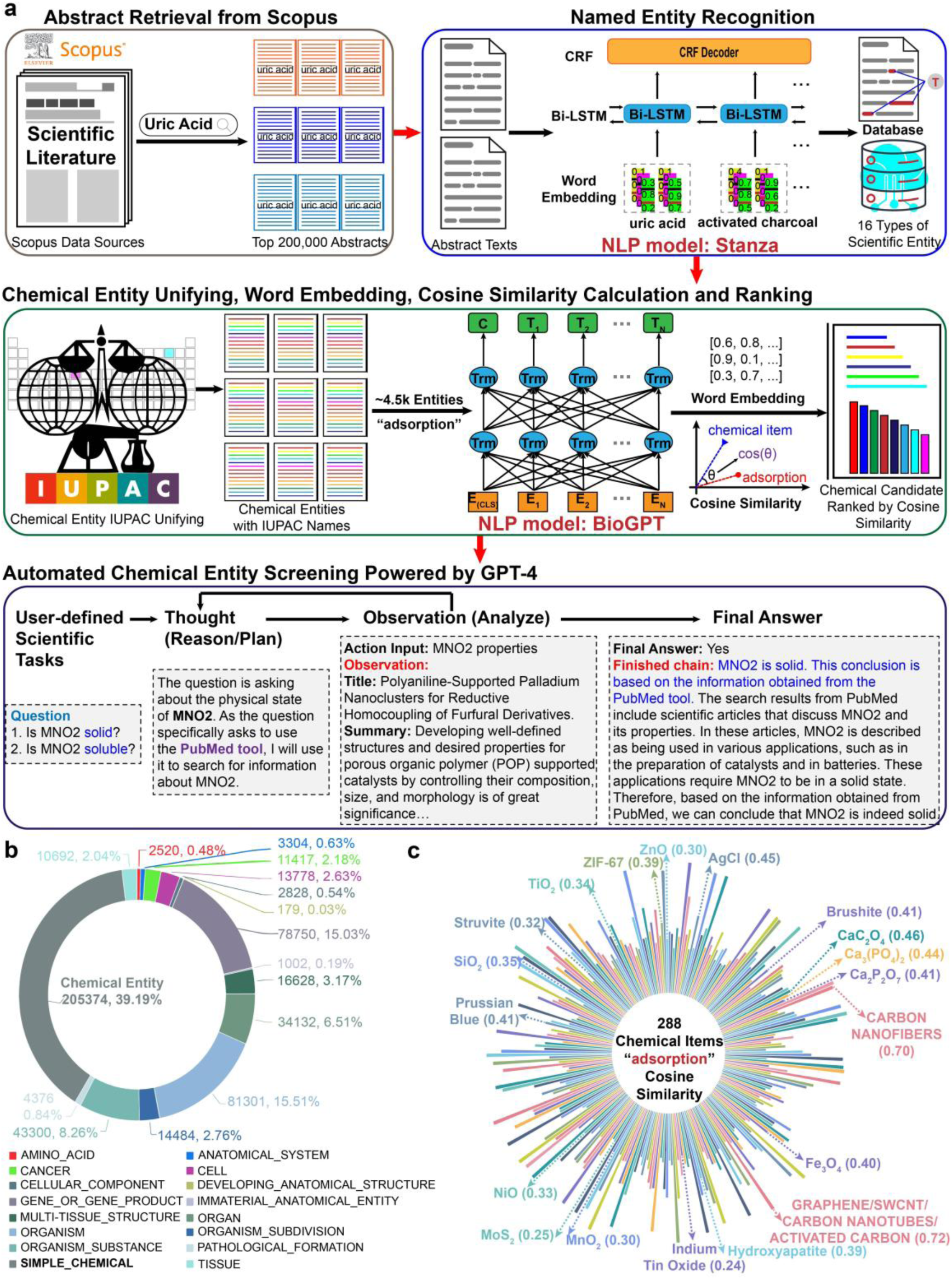
UA removal material discovery process. a) The schematic diagram for MatWISE (Material Workflow for Intelligent Screening and Evaluation) powered by NLP models. b) The names, counts and proportions of 16 types of scientific entity recognized from 20,000 abstracts of UA related literatures. The number of chemical entities amounted to 205374, accounting for 39.19% of the total entity (524065). c) Nightingale rose diagram for the cosine similarity scores (0.24 to 0.76) between the word embeddings of “adsorption” and 288 chemical entities from UA related literatures. The cosine similarity score could be leveraged to rank entities and ascertain those most plausibly affiliated with “uric acid adsorption”.

In recommendation systems, cosine similarity is commonly used to evaluate similarity between word embeddings of search results and search term.^[31]^ It is reasonable to assume that when similarity between the word embedding of a material from UA related literature and the embedding of ‘adsorption’ is high, the text corpus likely contains information concerning the adsorption behavior of the material (Figure S1a, Supporting Information, activated charcoal with 0.72 cosine similarity), whereas low similarity often denotes irrelevance (Figure S1c, Supporting Information, methyl cellulose with −0.01 cosine similarity and zero UA adsorption capacity). We analyzed the cosine similarities of the ∼4.5k chemical entities and identified that entities with cosine similarity below the average value were unrelated to our target keyword. Initial analysis revealed that entities with cosine similarities below the average value were unlikely to be related to UA adsorption. We set this average score as our threshold and selected entities that surpassed it. Our subsequent refinement involved considering the frequency of occurrence of each chemical entity in the database. Only those with frequencies higher than the average were retained, resulting in a shortlist of 288 IUPAC chemical entities (Figure 2c). These entities displayed clear clustering tendencies in three-dimensional space after undergoing t-distributed stochastic neighbor embedding (t-SNE)^[32]^ dimension reduction (Figure S1d, Supporting Information).

Subsequently, given numerous candidates with high potential of UA adsorption, we automatically eliminate those with undesired physical chemical properties using a tool-augmented large language model (LLM) powered by GPT-4 based on LangChain framework^[33]^ incorporating WebSearch^[34]^ and PubMedQuery. Test outcomes revealed consistent precision ranging from 97% to 99%, on evaluating insoluble and solid chemical attributes. With the feedback of physical chemical characteristics, we were able to filter 86% of materials not conforming to our desired characteristics.

At last, we selected 24 low-cost (less than $3 per gram) materials (Figure S1e, Supporting Information) after excluding absorbable organic solid compounds from the water insoluble solid materials to evaluate the UA adsorption ability. After primary screening, we got 6 materials with relatively good UA adsorption performance (∼40 mg g^-1^, Q_max_ = 42 mg g^-1^) (Figure S1e, Supporting Information). The top 1 material was identified after the second-round screening. MnO_2_ showed the highest adsorption capacity of 487 mg g^-1^ in simulated intestinal fluid (SIF) (Figure S1f, Supporting Information). Our integrated approach, combining MatWISE with stepwise experimental verifications suggests that MnO_2_ has the highest capacity to eliminate UA among all reported materials.

### 2.2. Synthesis and characterization of α, β, γ, δ-MnO_2_

To study the impact of crystal structure on UA adsorption, four major crystalline MnO_2_ including hollandite (α-MnO_2_), pyrolusite (β-MnO_2_), nsutite (γ-MnO_2_), and birnessite (δ-MnO_2_) were prepared by the one-step hydrothermal method in autoclaves (Figure S2a, Supporting Information). The XRD patterns (Figure 3a), X-ray photoelectron spectra (XPS, Figure 3b and Figure S2b, Supporting Information), Raman spectra (Figure S2c, Supporting Information), Fourier transform infrared (FTIR) analysis (Figure S2d, Supporting Information), energy dispersive spectrometer (EDS, Figure S2e-h, Supporting Information), scanning electron microscopy (SEM), and high-resolution (HR) transmission electron microscopy (TEM) images (Figure 3c-f) confirm the successful synthesis of different MnO_2_. As seen in Figure S2i, Supporting Information, δ-MnO_2_ has the most total weight loss (19.8%), namely water content, followed by γ-MnO_2_ > β-MnO_2_ > α-MnO_2_. Finally, the oxygen vacancy content of these samples was evaluated by electron paramagnetic resonance (EPR), following the order of δ-MnO_2_ > γ-MnO_2_ > β-MnO_2_ > α-MnO_2_ (Figure S2j, Supporting Information).

**Figure 3.**
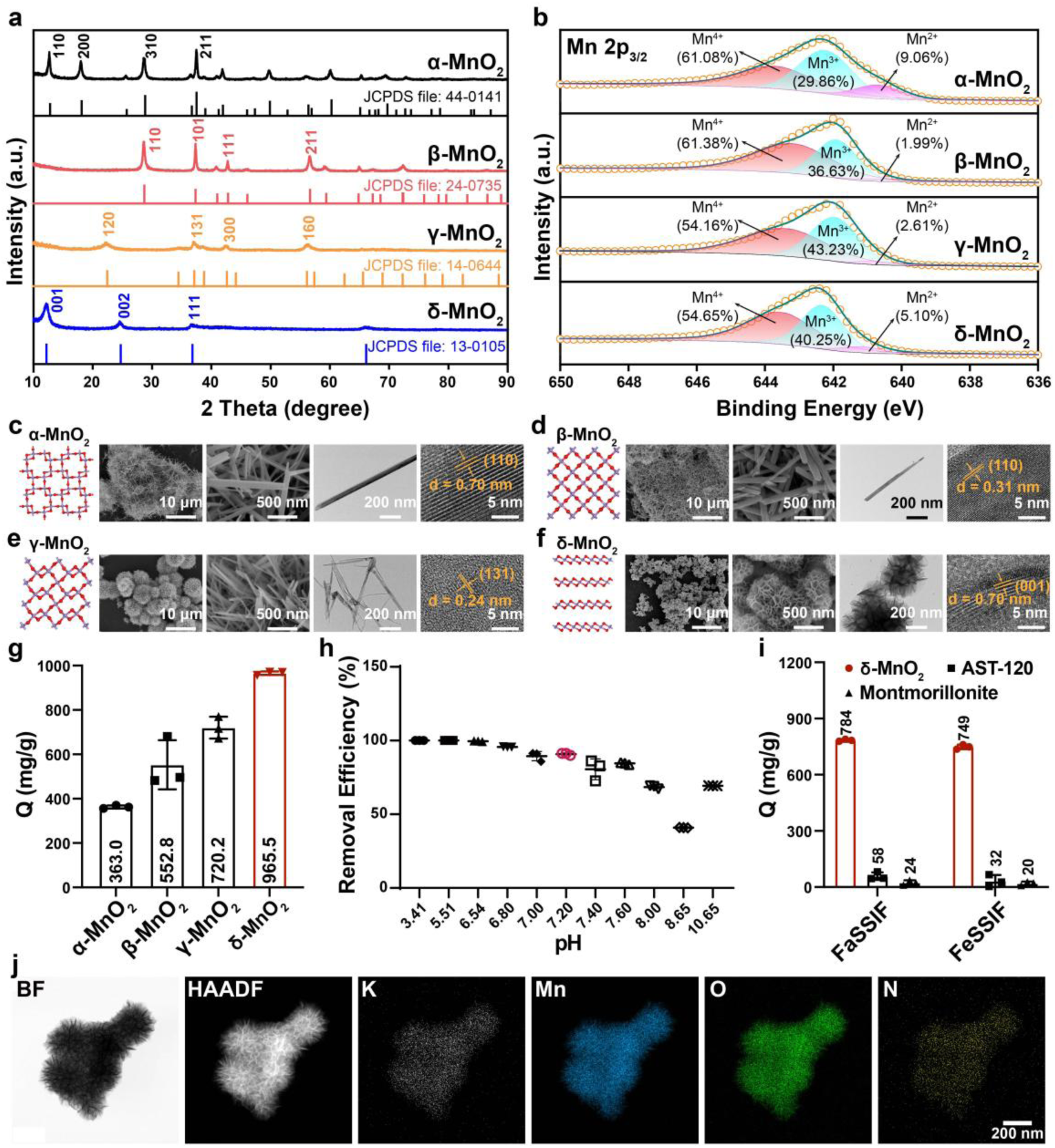
Characterizations and UA adsorption study of MnO_2_. a,b) The XRD (a) and Mn 2p_3/2_ XPS (b) spectra of α, β, γ, δ-MnO_2_. c-f) Schematic diagrams of crystal structure (red: O, purple: Mn), SEM (scale bars: 10 μm or 500 nm), TEM (scale bars: 200 nm), and HRTEM (scale bars: 5 nm) images of MnO_2_. g) Adsorption capacities of UA onto MnO_2_ in SIF (n = 3 replicates). h) Effect of solution pH on UA adsorption onto δ-MnO_2_ (n = 3 replicates). i) Adsorption capacities of δ-MnO_2_, AST-120, and montmorillonite regarding UA adsorption in FaSSIF or FeSSIF (n = 3 replicates). j) FETEM-EDS mapping of BF, HAADF images and elemental mapping images of K, Mn, O, and N for δ-MnO_2_ adsorbed UA sample. Scale bar: 200 nm.

### 2.3. Adsorption property of UA onto MnO_2_

To explore the UA adsorption capacities of four types of MnO_2_, we conducted the *in vitro* adsorption experiment in SIF. The adsorption capacities are followed the order of δ-MnO_2_ > γ-MnO_2_ > β-MnO_2_ > α-MnO_2_ (Figure 3g), which strongly correlate to the content of water (Figure S2i, Supporting Information) and oxygen vacancies (Figure S2j, Supporting Information) in MnO_2_, but not to the amounts of Mn^2+^, Mn^3+^, Mn^4+^, O_ad_, and O_la_ (Figure 3b and Figure S2b, Supporting Information), the amounts of BET specific surface area, total pore volume or pore size (Figure S2k-m, Supporting Information). Density function theory (DFT) calculation also showed that δ-MnO_2_ has the highest adsorption energy than that of another three MnO_2_ (Figure S3a-d, Supporting Information). The central composite design of response surface methodology (RSM-CCD, Figure S4, Supporting Information), Pearson’s correlation coefficient (PCC, Figure S5a, Supporting Information) and machine learning (ML, Figure S5b-r, Supporting Information) were used to investigate the optimization and interactions of experimental factors of UA adsorption onto δ-MnO_2_, implying that the solution pH is the most important factor affecting adsorption. δ-MnO_2_ has a strong removal ability of UA at the physiological pH of intestine (pH 7.2, Figure 3h), where UA mostly presents.

Subsequently, kinetic (Figure S6a-d, Supporting Information) and isotherm (Figure S6e-h, Supporting Information) study showed that the adsorption process is a chemical sorption, and the maximum adsorption capacity (Q_m_) is 1357.22 mg g^-1^, which is the highest among all reported UA adsorbents.^[35–41]^ Finally, thermodynamic study (Figure S6i-o, Supporting Information) exhibited that the adsorption process is spontaneous, heat-absorbing, and irreversible due to the negative standard free energy change (ΔG^0^), positive standard enthalpy (ΔH^0^) and entropy (ΔS^0^) changes.

### 2.4. Adsorption selectivity and oxidation mechanism

We investigated how inorganic and organic compounds in the GI tract affect the UA adsorption onto δ-MnO_2_. There is no obvious impact on the adsorption capacity in the presence of inorganic salt and most organic nutrients except sodium oleate and ascorbic acid (Figure S3e-g, Supporting Information), which have limited concentrations in the GI tract. The high selectivity coefficient (Figure S3h, Supporting Information) and almost zero UA desorption (Figure S3i, Supporting Information) reveal the extraordinarily high recognition selectivity and binding affinity of UA onto δ-MnO_2_. Furthermore, δ-MnO_2_ retains a very high adsorption capacity in fasted-state simulated intestinal fluid (FaSSIF) and fed-state simulated intestinal fluid (FeSSIF), which is 15-40 times greater than that of AST-120 and montmorillonite (Figure 3i). EDS mapping, FTIR, XPS, and Raman were conducted to understand the adsorption mechanism of UA onto δ-MnO_2_ (Figure S3j-m, Supporting Information). XPS analysis revealed that there is a redox reaction between UA and δ-MnO_2_ since the proportion of Mn^2+^ increased (from 5.10% to 15.16%) and O decreased (from 30.59% to 27.95%) after adsorption of UA (Figure 3b, Figure S2b, S3k-l, Supporting Information). Using inductively coupled plasma optical emission spectrometry (ICP-OES), magnetic resonance imaging (MRI) and liquid chromatography-mass spectrometry (LC-MS) analyses on the solution after adsorption (Figure S3n-q, Supporting Information), we further validated the increased Mn^2+^ content and the formation of oxidation product allantoin (Figure S3r, Supporting Information) after adsorption. These findings suggest that δ-MnO_2_ has a significant potential to remove UA in the GI tract by dual adsorption and oxidation.

### 2.5. Toxicity assessment

We performed *in vitro* Mn release and *in vivo* mouse experiments to assess the toxicity of δ-MnO_2_. There are no detectable manganese ions release from δ-MnO_2_ in simulated gastric fluid (SGF) and SIF (Figure 4a). The insolubility nature of δ-MnO_2_ should guarantee the safety when orally administered. In the mice study, δ-MnO_2_ only distributed in and quickly passed through GI tract after oral administration (Figure 4b-c). δ-MnO_2_ was entirely eliminated from the GI system within 8 hours (Figure 4d), suggesting that MnO_2_ does not adhere to GI tract. We further evaluated the various health indicators and manganese uptake of mice after repeated oral administration for two weeks (Figure 4e). No side effects were observed in C57BL/6J mice after oral administration of δ-MnO_2_ at four different high doses (0.25, 0.5, 1, 2 g/kg/day). There was no significant change in body weight (Figure S7a, Supporting Information), serum biochemical analysis (Figure S7b-i, Supporting Information) for liver and kidney functions, blood routine test (Figure S7j-y, Supporting Information) among different doses of δ-MnO_2_ and the control group. There was no discernible change in the manganese level in their major organs, blood cells, and hair (Figure 4f) after two weeks’ oral administration, except a minimal manganese level increase in liver, which returned to base level followed by a 7-day recovery (Figure 4f, inserted). Moreover, the histological images of major organs in various dosages of δ-MnO_2_ groups showed no pathological changes from the control group (Figure 4g and Figure S8, Supporting Information). It is worth mentioning that MnO_2_ does not increase the burden on the kidneys as Mn has a minimal elimination through and no accumulation in the kidney.^[42]^ These findings together demonstrate the non-toxic nature of orally administrated δ-MnO_2_, and especially suitable for renally impaired patients.

**Figure 4.**
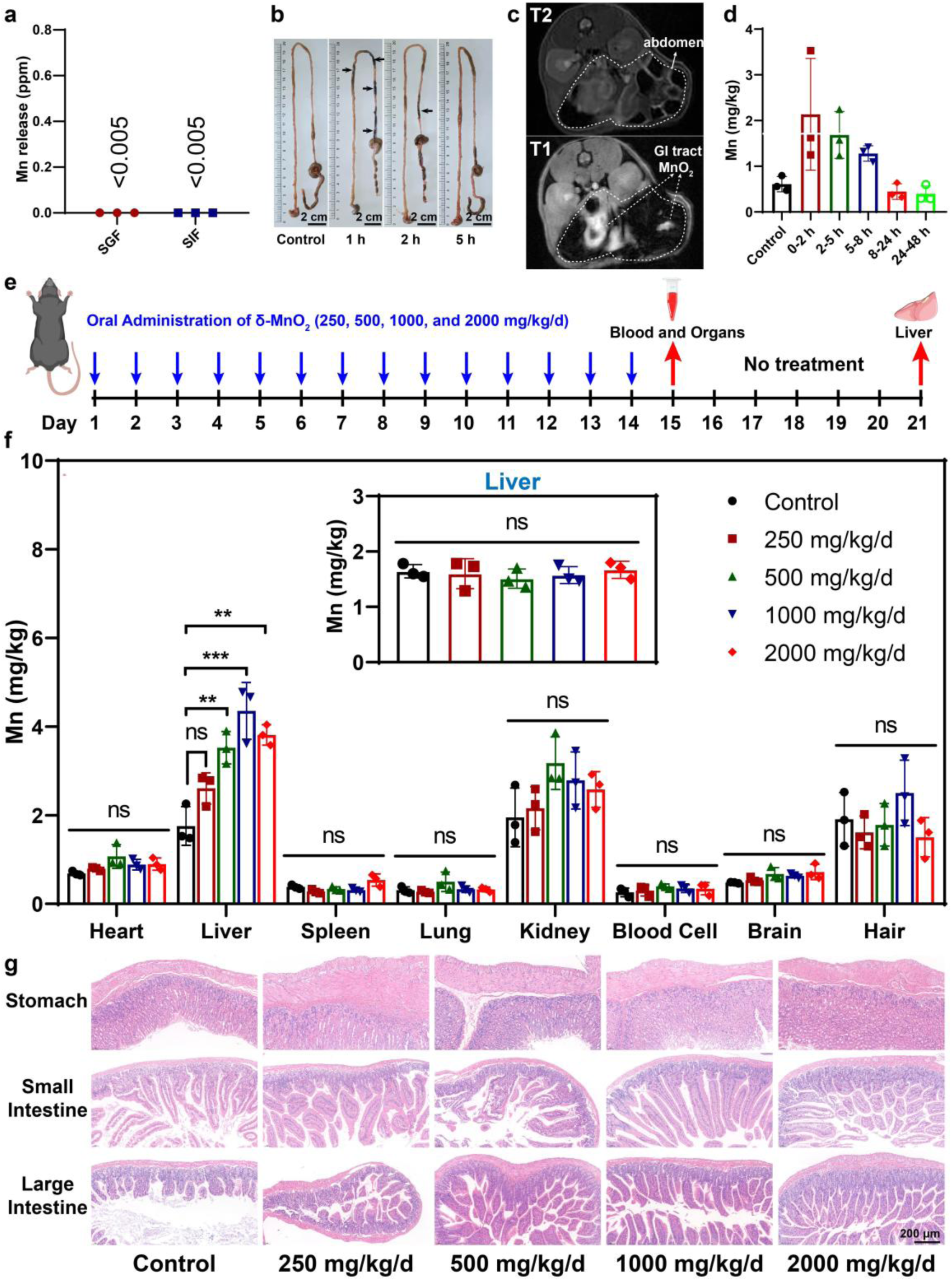
Biosafety assessment of orally administrated δ-MnO_2_. a) The manganese release from δ-MnO_2_ in SGF and SIF at 37 ℃ for 4 h. b) Representative photographs of GI tract in mice at 0, 1, 2, and 5 h after gavage administration of δ-MnO_2_ at a dose of 250 mg/kg. The arrow points MnO_2_. Scale bar: 2 cm. c) The MRI images of mice at 1 h after oral administration of δ-MnO_2_. d) The manganese content in the feces excreted from C57BL/6J mice after oral administration of δ-MnO_2_ (n = 3). e) Treatment schedule of *in vivo* biosafety assessment for δ-MnO_2_. f) Manganese distribution in the organs and tissues of C57BL/6J mice after 14-day repeated oral dosing of δ-MnO_2_ (n = 3). Insert graph depicted the manganese level in livers of mice treated with δ-MnO_2_ for 14 days and followed by a 7-day pause. ns, not significant; **p<0.01; ***p<0.001, ordinary one-way analysis of variance (ANOVA). g) Representative hematoxylin and eosin (H&E) staining images of stomach, small and large intestine in mice treated with δ-MnO_2_ for 14 days. Scale bar: 200 μm.

2.6. *In vivo* UA-lowering experiment

We employed three different hyperuricemia mouse models to thoroughly study the *in vivo* UA-lowering effect of δ-MnO_2_. First, a stable short-term hyperuricemia model was established by intraperitoneal injection of potassium oxonate (PO) combined with intragastric administration of UA to mice (Figure 5a and 5b). The blood UA was significantly lowered after oral administration of δ-MnO_2_ (Figure 5b).

**Figure 5.**
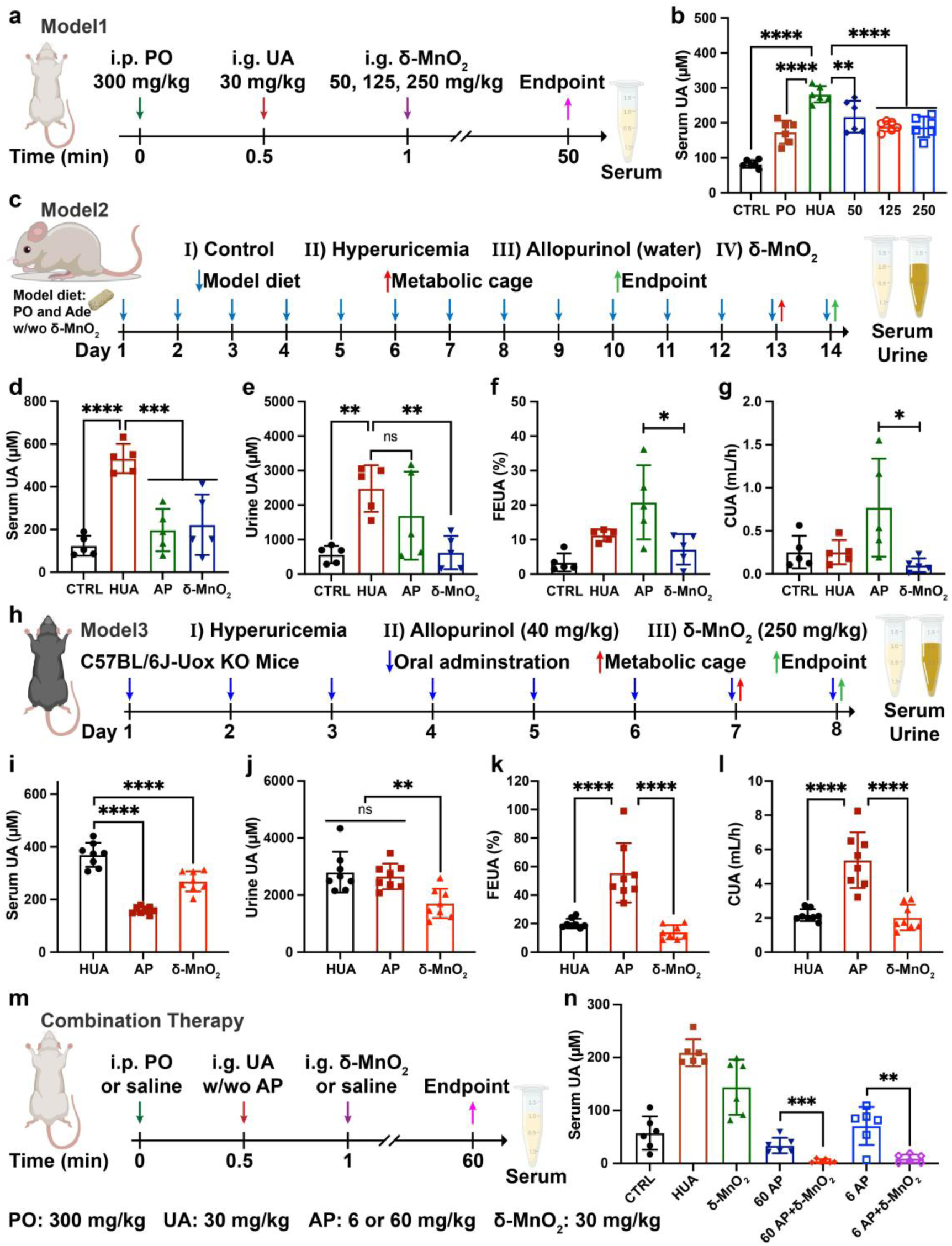
Oral administration of δ-MnO_2_ lowering UA in three mouse models. a) Schedule for short-term UA-lowering experiment. b) Serum UA levels in the normal and hyperuricemia mice with or without administration of δ-MnO_2_ (n = 6). Control (CTRL), Hyperuricemia (HUA) and δ-MnO_2_ (50, 125, and 250 mg/kg). c) Schedule of long-term UA-lowering animal experiment. Allopurinol (AP). d-g) Serum UA (d), urinary UA (e), FEUA (f) and CUA (g) in PO and Ade induced hyperuricemia mice (n = 5). h) Schedule of 7-day UA-lowering animal experiment using C57BL/6J-Uox KO mice. i-l) Serum UA (i), urinary UA (j), FEUA (k), and CUA (l) in Uox KO mice (n = 8). m) Schedule for combination therapy of δ-MnO_2_ and AP for UA-lowering treatment. n) The serum UA levels in the normal and hyperuricemia mice of combinative administration of δ-MnO_2_ and AP (n = 6). Data are shown as mean ±SD. ns, not significant; *p<0.05; **p<0.01; ***p<0.001; ****p<0.0001, ordinary one-way analysis of variance (ANOVA, Figure 5b, d-e, and i-l) or two-tailed unpaired t-test (Figure 5f, g, and n).

As the second model, we established a long-term hyperuricemia mouse model fed with a diet containing PO and adenine (Ade) for 14 days (Figure 5c) and used the clinical standard-of-care allopurinol as the positive control. The serum and urine UA levels of the model group were significantly higher than those of the normal mice (Figure 5d and 5e). After the treatment of δ-MnO_2_ with feed or allopurinol drinking water during the modeling process, the blood UA level significantly decreased (Figure 5d). The urine UA level in the δ-MnO_2_ group was significantly lower than the model group and close to the normal mice, whereas allopurinol treatment did not significantly decrease urine UA (Figure 5e). The excretion fraction of uric acid (FEUA) and clearance rate of uric acid (CUA) were significantly lower in the δ-MnO_2_ group compared to the group treated with allopurinol (Figure 5f and 5g).

Moreover, we evaluated the UA-lowering capability of δ-MnO_2_ in a spontaneous hyperuricemia mouse model (C57BL/6J-Uox KO mice), which is a more suitable model mimicking the human UA disorders (Figure 5h). As shown in Figure 5i-l, the serum and urine UA levels in the δ-MnO_2_ group were significantly lower than those in the hyperuricemia model, as well as the FEUA and CUA were significantly lower than those in the allopurinol group. δ-MnO_2_ could reduce the amount of UA excreted through kidney and ease the burden of kidney, which is important to protect residual renal functions of renally impaired hyperuricemia patients. δ-MnO_2_ may provide a long-waited clinical solution for CKD patients who have no suitable UA-lowering drugs while protecting of residual renal functions.^[16]^

Additionally, the combination of δ-MnO_2_ and allopurinol can significantly reduce the blood UA even with even one-tenth of the lowest recommended dose of allopurinol (Figure 5m-n), thanks to the complementary dual effects of inhibiting production and promoting intestinal elimination. This study implies that combination therapy may reduce side effects of allopurinol by significantly lowering its dose. Our findings suggest that the orally administered, nonabsorbable δ-MnO_2_ may be more clinically beneficial than currently available UA-lowering medications, or at least clinically meaningful when used as combo, by providing a new route to reduce serum UA level, especially for individuals with impaired kidney functions not suitable for the standard-of-care anti-hyperuricemia medications.^[43]^

## 3. Conclusion

For the treatment hyperuricemia, we designed a multi-tiered framework MatWISE integrated with advanced AI techniques to tap the latent knowledge embedded within extant research corpora in vast amounts of literature in search of materials with the potential to absorb UA. Through the experiment screening, MnO_2_ was identified as the most promising starting point. In the next step optimization, δ-MnO_2_ emerged with an unprecedent high UA adsorption capacity (absorbing nearly the equivalent weight of UA *in vitro*). Looking back to the literatures, MnO_2_ was only reported as a detection reagent for UA in analytical chemistry,^[44]^ and the most valuable δ-MnO_2_ was never reported in literatures for any interactions with UA.

δ-MnO_2_ enables the rapid and highly selective removal of UA even in the complex GI tract environment. δ-MnO_2_ has the following advantages: (i) excellent stability and insoluble in the whole GI tract; (ii) fast adsorption kinetics and high adsorption capacity; (iii) no desorption; (iv) negligible manganese uptake, almost no excretion through kidney and non-toxic *in vivo*; (v) effectively lower blood UA and urine UA, and the ability of reducing urine UA better than that of the commercially available allopurinol in the hyperuricemia mouse models. Furthermore, the combination of δ-MnO_2_ and clinical XOIs not only achieves normal serum UA levels but also significantly reduces the dosage of both drugs. Overall, these results suggest that oral administration of δ-MnO_2_ has great clinical potential, which may provide a novel and kidney-friendly therapeutic option for hyperuricemia patients, and reduce the renal UA burden in chronic kidney disease patients with hyperuricemia. This proof-of-concept study for UA-lowering biomaterial discovery using a seamless integration of BioGPT and GPT-4 empowered MatWISE, and a handful of experimental screening efficiently discover a translationally relevant material for an unmet clinical need.

We hope that this work will inspire a new research paradigm for leveraging AI to fully utilize the latent knowledge embedded within existing research corpora to facilitate the translation of more scientific findings into practical applications. This new research paradigm enables the fair selection and comparison among all possible solutions by unfolding latent knowledge in literatures. And more importantly, it may have the emergent ability that triggers the discovery of materials with unprecedent functions in the uncharted space.

## 4. Experimental Section/Methods

### Materials and Methods

All materials, methods, supplementary references, and datasets are available in **Supporting Information**. All the animal experiments were conducted following the protocols approved by the Institutional Animal Care and Use Committee of Shanghai Jiao Tong University (No. A2022078).

### Statistical analysis

All data were reported as mean ±standard deviation (SD). Statistical analysis was conducted using GraphPad Prism version 8.0. A Student’s t-test or a one-way ANOVA with tukey’s multiple comparisons test was performed when comparing two groups or more than two groups, respectively. Difference was statistically significant if p < 0.05.

## Supporting Information

Supporting Information is available from the Wiley Online Library or from the author.

## Acknowledgements

We would like to thank X. Liu for help in collecting MRI images. This work also supported by the grants from Interdisciplinary Program of Shanghai Jiao Tong University (YG2022ZD002), the National Natural Science Foundation of China (Grant No. 22171184), and China Postdoctoral Science Foundation (2022M712091).

## Author Contributions

X.D.Z., X.Z., J.L. and S.Z. conceived and led the research. J.Q. and J.A. developed the data processing pipeline MatWISE and analysed the literature mining results. J.L. and M.W. supervised the establishment of the pipeline. X.D.Z., K.L., L.X. and T.L. performed material preparation, characterization and adsorption study. X.D.Z. and X.Z. performed UA-lowering experiment and toxicity assessment. X.D.Z., J.L. and S.Z. wrote the initial manuscript with input from J.Q., K.L. and J.A. All authors contributed to reviewing and editing the manuscript.

## Data Availability Statement

The data that support the findings of this study are available from the corresponding author upon reasonable request. We release an open-source implementation of MatWISE at https://github.com/Junhang0202/MatWISE. Beyond the methods described so far, this release includes simple demo and installation guide to help users generating results that are reported in the paper.

Received: ((will be filled in by the editorial staff))

Revised: ((will be filled in by the editorial staff))

Published online: ((will be filled in by the editorial staff))

---

A multi-tiered framework, MatWISE, that integrates natural language processing, semantic relationship mapping, and machine learning to automate the complex process of material discovery from vast scientific literatures is designed. It successfully identifies δ-MnO_2_ for efficiently lowering serum uric acid levels in three hyperuricemia models via oral administration, demonstrating the feasibility of developing new drugs assisted by artificial intelligence.

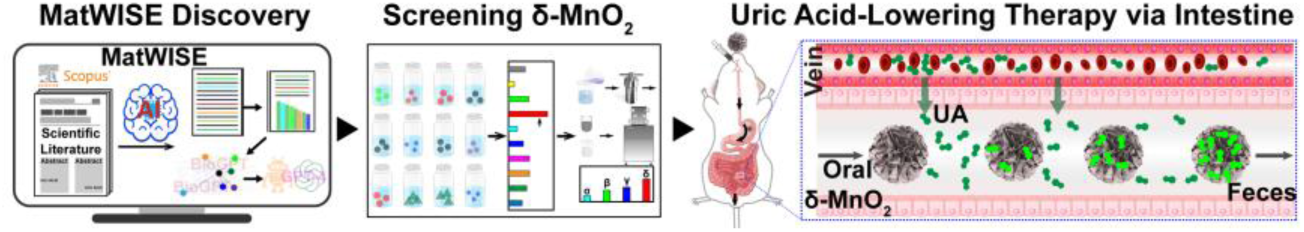

