## Supporting Information for "Automated Discovery of Therapeutic Biomaterial for Renally Impaired Hyperuricemia Patients by Natural Language Processing and Machine Learning"

### Materials and Methods

#### Materials

The chemical reagents used in experiments include potassium permanganate ($KMnO_4$), manganese sulfate monohydrate ($MnSO_4 \cdot H_2O$), ammonium persulfate ($(NH_4)_2S_2O_8$), potassium dihydrogen phosphate ($KH_2PO_4$), sodium hydroxide (NaOH), potassium oxonate, uric acid, allopurinol, adenine, and so on. All of these reagents were purchased from Energy Chemical, Aladdin, and Tansoole (China) or Sigma-Aldrich (USA). These were all analytical grade chemicals and used directly without further purification. One liter of simulated intestinal fluid (SIF, pH 6.8) was made by dissolving 6.8 g of $KH_2PO_4$ in distilled deionized water, elevating pH to 6.8 using 0.5 M NaOH aqueous solution and adjusting to 1000 mL with distilled deionized water. One liter of simulated gastric fluid (SGF, pH 1.2) was made by dissolving sodium chloride (2 g) in distilled deionized water, adjusting pH to 1.2 using 1 M HCl aqueous solution, and then adjusting the volume to 1000 mL with distilled deionized water. One liter of fasted-state simulated intestinal fluid (FaSSIF, pH 6.5) was obtained by dissolving sodium taurocholate (1.613 g), maleic acid (2.219 g), sodium hydroxide (1.392 g), sodium chloride (4.014 g) and lecithin (0.152 g) in 1000 mL distilled deionized water. One liter of fed-state simulated intestinal fluid (FeSSIF, pH 5.8) was prepared by dissolving sodium taurocholate (5.377 g), sodium oleate (0.244 g), maleic acid (6.386 g), sodium hydroxide (3.266 g), sodium chloride (7.342 g), lecithin (1.516 g), and glyceryl monooleate (1.782 g) in 1000 mL distilled deionized water.

#### Discovering UA related materials using NLP techniques

We designed an automated system dubbed MatWISE for streamlining the search and selection procedure of UA related materials form literatures. The codes are available at https://github.com/Junhang0202/MatWISE. The keywords "uric acid" was used to retrieve literatures in Scopus (https://www.scopus.com/) database and extract their titles and abstracts from the first 20,000 articles sorted by relevance. To identify chemical named entities recognition, a specialized extension of the Stanza library, specifically the biomedical and clinical English model packages, was employed. This NLP model was trained using three corpuses related to chemical compounds: BC5CDR[1], BC4CHEMD[2], and BioNLP13CG[3].

According to our experiment, the Stanza model trained with BioNLP13CG corpus was selected due to its superior performance and comprehensive coverage of 16 distinct entity types.

Subsequently, to further enhance the accuracy of analyzing chemical entities, we employed a systematic approach to handle the non-standard chemical entity names in the dataset. More specifically, we converted these non-standard chemical entity names to a systematic naming convention provided by the International Union of Pure and Applied Chemistry (IUPAC) nomenclature. Furthermore, a pretrained domain-specific generative Transformer language model BioGPT was utilized to convert each chemical entity into a word embedding vector. Word embedding is a powerful technique that assigns a vector representation to each word in a given article. They capture the semantic and syntactic relationships between words, allowing the NLP model to understand the meaning and context of the text more effectively. These vectors are positioned in a high-dimensional space in which words with similar meanings are closer to each other, while words with different meanings are further apart. This proximity is measured in terms of cosine similarity, which is a common approach in recommendation systems to measure the similarity between two vectors. Words with similar semantics in the vector space tend to have higher cosine similarity, while words with dissimilar semantics have lower cosine similarity.

To identify chemical entities relevant to UA adsorption, we employed a two-step approach. Firstly, we calculated the cosine similarity between each entity and the keyword "adsorption" to measure their semantic similarity. This allowed us to obtain similarity scores, which served as the basis for ranking the entities with the domain of "uric acid adsorption". In addition to cosine similarity calculations, we utilized an Exploratory Data Analysis (EDA) process for the recommendation. Initially, we analyzed the cosine similarities of approximately 4.5k chemical entities and identified those with similarity scores below the average value as unrelated to our target keyword. This enabled us to establish the average cosine similarity score as a threshold and select entities with similarity scores surpassing this threshold as candidates. Furthermore, we refined the candidate list by considering the occurrence frequency of the chemical entities in the database, retaining entities with frequencies higher than the average occurrence frequency value. By combining these methods, we aimed to identify chemically relevant candidates for UA adsorption. To gain insights hidden relationships within the chemical entities, we employed a data reduction method called t-distributed stochastic neighbor embedding (t-SNE) algorithm. This technique reduced the dimensionality of the word embeddings of the candidate chemical entities, enabling us to visualize the data in a three-dimensional space. By plotting the distribution maps derived from the dimensionality reduction data using t-SNE visualization, we were able to identify patterns and clusters that may indicate potential relationships between the chemical entities.

Considering our objective of screening and removing UA material from the gastrointestinal tract, we also aim to eliminate small organic molecules and water-soluble inorganic compounds that can be absorbed by the gastrointestinal tract. Drawing inspiration from successful applications in other fields[4], we have integrated MatWISE and GPT-4, to simplify the compound screening process. In our experiment, MatWISE primarily utilizes two tools, WebSearch and PubMedQuery, to guide GPT-4 in accomplishing specific tasks. Specifically, MatWISE enables GPT-4 to perform tasks by invoking different tools. When using MatWISE, a list of tool names, descriptions of their purposes, as well as detailed information about expected inputs and outputs are required. Subsequently, guided by user prompts, GPT-4, serving as the core of MatWISE, utilizes the provided tools when necessary to complete the tasks. In this research, we employed the use of MatWISE to conduct experiments aimed at answering two crucial questions regarding the current compound's solubility and physical state in water, namely whether it exists in a solid state or not. Following the screening facilitated by MatWISE, the assessment of a chemical expert was sought to evaluate the effectiveness in addressing the aforementioned issues. Experimental results show that MatWISE is highly accurate in handling these chemical queries, with accuracy rates ranging from 97% to 99%. We used MatWISE in our experiments to automatically filter the remaining compound entities, thus improving the overall efficiency of the screening.

**Batch experiments for UA adsorption screening**

A typical UA adsorption experiment of adsorbents was carried out in a 50 mL plastic centrifuge tube containing an aqueous SIF solution of UA (500 μM, $C_0$) and adsorbent powder (0.1 or 2 g $L^{-1}$, m/V). The mixture was shaken in a shaker for 4 h at 37 °C and then filtered by a polyethersulfone (PES, 0.22 μm) filter for UA measurement. The final UA concentration (C) of the filtrate was detected using a microplate reader at 291 nm.

The adsorption capacity (Q, mg $g^{-1}$) was calculated using the following equation:

$$Q = 0.168 * (C_0 - C) * \frac{V}{m}$$

where $C_0$ and C are the UA concentration (μM) at the initial solution and after adsorption, respectively, V is the volume (mL) of the solution, and m is the mass (mg) of adsorbents.

**Synthesis of α, β, γ, δ-$MnO_2$**

Synthesis of α-$MnO_2$. 0.625 g $KMnO_4$ and 0.265 g $MnSO_4·H_2O$ were dissolved in 40 mL deionized water and stirred at room temperature for 30 min to form a homogeneous solution. Then, the mixture was transferred to a 100 ml Teflon-lined stainless-steel autoclave and heated in a drying oven at 160 ℃ for 12 h. After cooling to room temperature naturally, the solids were collected by filtration on a sand core funnel, washed with deionized water for 5 times, and dried overnight by freeze-dryer to obtain the product with the yield of 98.2%.

Synthesis of β-$MnO_2$. 0.845 g $MnSO_4·H_2O$ and 1.14 g $(NH_4)_2S_2O_8$ were dissolved in 40 mL deionized water and stirred at room temperature for 5 min to form a homogeneous solution. Then, the mixture was transferred to a 100 ml Teflon-lined stainless-steel autoclave and heated in a drying oven at 140 ℃ for 12 h. After cooling to room temperature naturally, the solids were collected by filtration on a sand core funnel, washed with deionized water for 5 times, and dried overnight by freeze-dryer to obtain the product with the yield of 72.6%.

Synthesis of γ-$MnO_2$. 1.69 g $MnSO_4·H_2O$ and 2.29 g $(NH_4)_2S_2O_8$ were dissolved in 40 mL deionized water and stirred at room temperature for 10 min to form a homogeneous solution. Then, the mixture was transferred to a 100 ml Teflon-lined stainless-steel autoclave and heated in a drying oven at 90 ℃ for 24 h. After cooling to room temperature naturally, the solids were collected by filtration on a sand core funnel, washed with deionized water for 5 times, and dried overnight by freeze-dryer to obtain the product with the yield of 70.4%.

Synthesis of δ-$MnO_2$. 0.75 g $KMnO_4$ and 0.14 g $MnSO_4·H_2O$ were dissolved in 40 mL deionized water and stirred at room temperature for 30 min to form a homogeneous solution. Then, the mixture was transferred to a 100 ml Teflon-lined stainless-steel autoclave and heated in a drying oven at 160 ℃ for 12 h. After cooling to room temperature naturally, the solids were collected by filtration on a sand core funnel, washed with deionized water for 5 times, and dried overnight by freeze-dryer to obtain the product with the yield of 99.3%.

**Characterizations of α, β, γ, δ-$MnO_2$**

Various analytical methods were used to characterize $MnO_2$. Powder X-ray diffraction (XRD) was used to investigate the crystal structure of $MnO_2$ using a Bruker D8 Advance X-ray diffractometer operating in a Da Vinci geometry equipped with a Lynxeye detector and a radioactive Cu-Kα source. Particle morphologies were investigated using scanning electron microscopy (SEM, RISE-MAGNA, TESCAN). The quantitative analysis and mapping of the elements (Mn, O, K, or S) of $MnO_2$ were obtained using energy dispersive X-ray spectroscopy (EDS, RISE-MAGNA, TESCAN). A field-emission transmission electron microscopy (FETEM, Talos F200X G2) was used to examine the high-resolution transmission electron microscopy (HRTEM) micrographs of $MnO_2$ or EDS mapping of the elements (Mn, O, K, and N) for the samples of UA adsorbed onto δ-$MnO_2$. X-ray photoelectron spectroscopy (XPS) analysis were utilized to investigate the surface elemental states of samples using a Kratos Axis Ultra DLD spectrometer with a monochromated Al-Kα X-ray source. The Brunauer-Emmett-Teller (BET) surface area, pore size, and pore volume of $MnO_2$ were characterized by nitrogen adsorption-desorption at liquid nitrogen temperature using the Quantachrome Autosorb iQ3 instrument. The thermal stability and thermal degradation of $MnO_2$ were investigated using thermogravimetry (TGA) in a nitrogen atmosphere utilizing a NETZSCH STA 449F3 thermo gravimetric analyzer with a heating rate of 10 °C $min^{-1}$ from ambient temperature to 900 °C. $MnO_2$ or UA powders were combined with KBr for Fourier transformed infrared (FTIR) measurement on a Nicolet 6700 spectrometer between 400 and 4000 $cm^{-1}$. Raman spectra were acquired using a confocal Raman microscope system (Renishaw inVia Qontor, UK). Electron paramagnetic resonance (EPR) spectra were collected on a Bruker A300 X-band spectrometer at 100 K.

**Optimization of adsorption condition**

*Design of experiments using response surface methodology (RSM)*

In this study, we introduced a mathematical and statistical modeling technique, RSM with central composite design (CCD), for optimization of critical adsorption parameters of UA onto δ-$MnO_2$. The four factors used as independent variables were adsorbent dose, pH, contact time, and initial concentration. These factors altered at five different levels (-α, -1, 0, +1, +α, α=2), as indicated in Figure S9a. The adsorption capacity of UA was the dependent variable. Design of experiment, regression analysis, analysis of variance (ANOVA), and variable optimization of critical adsorption parameters for the UA adsorption process were all done with Design Expert software (version 13.0.1.0). As shown in Figure S9b, a total of 30 experimental data sets were used for RSM analysis. The acceptability of RSM model was determined using the regression coefficient ($R^2$) and the p-value from ANOVA. The validated model was used to generate a surface response in order to determine the best adsorption conditions.

The UA adsorption process in SIF solution was operated according to the information in Table S2. After the adsorption completion, the suspension was filtered through a 0.22 μm filter. The UA concentration in the filtrate was measured with a microplate reader at 291 nm. The adsorption capacity (Q, mg $g^{-1}$) was calculated using the following equation:

$$Q = 0.168 * \frac{(C_0 - C)}{Ad}$$

where $C_0$ and C are the initial concentration of UA (μM) and concentration after adsorption, respectively. Ad is the adsorbent dose (g $L^{-1}$).

*Machine learning (ML) algorithms*

Before ML analysis, the correlation between a set of input features for 176 data points was examined using Pearson's correlation coefficient (PCC). As shown in Figure S5a, the PCC value in the PCC matrix for pH and adsorption capacity was -0.81, indicating a strong negative correlation between these variables. There was no significant correlation (PCC values between -0.5 and 0.5) of the variables after excluding the pH and adsorption capacity, which helps to retain all these features for the construction of the ML predictive model. Subsequently, we employed decision trees, support vector machines (SVM), BP neural networks, extra trees, AdaBoost, XGBoost, random forests, GBDT, and CatBoost to predict adsorption capability and investigate the relationship between adsorption circumstances and adsorption capability of UA. The adsorbent dose, pH, contact time, ionic concentration (the concentration of potassium phosphate monobasic aqueous solution), temperature, initial UA concentration, and adsorption capacity were among the seven columns of data in this dataset, which had a total of 176 instances. The Scientific Platform Serving for Statistics Professional (SPSSPRO) was used to make all machine learning predictions and Pearson's correlation coefficient analysis. Adsorbent dosage, pH, contact time, ionic concentration, temperature, and initial UA concentration were the input variables. The UA adsorption capacity was the input response value. All algorithms were tested using 5-fold cross validation, which is a useful method for preventing data overfitting. The data was randomly divided into five groups, four for training and one for testing, and the procedure was done five times, each time with different data for testing. The reliability, conformity, and accuracy of the machine learning model were assessed using the correlation coefficient ($R^2$), mean absolute error (MAE), and root mean square error (RMSE), mean squared error (MSE), mean absolute percentage error (MAPE), which were all produced automatically by the SPSSPRO.

**UA removal capability in simulated GI fluids**

500 μM SIF, FaSSIF or FeSSIF solutions of UA were prepared. Adsorption experiments were carried out in 50 mL plastic centrifuge tubes containing 4 mg adsorbents and 40 mL UA solutions. The tubes were shaken at 180 rpm in a 37 °C constant temperature shaker for 4 h.

The sample was centrifuged at 10000 rpm for 5 min and the supernatant was filtered through a 0.22 μm filter. The absorbance of the filtrate at 291 nm was detected using a microplate reader. The UA standard solutions with different concentrations were prepared using the corresponding SIF, FaSSIF or FeSSIF solutions and followed by the same treatment with the adsorbent-treated samples. The UA concentration in the sample was calculated according to the standard curve. The percentage of UA removal (R%) was calculated using the following equation:

$$R\ (\%) = 100 * \frac{C_0 - C_e}{C_0}$$

The adsorption capacity ($Q_e$, mg $g^{-1}$) was calculated using the following equation:

$$Q_e = 0.168 * (C_0 - C_e) * \frac{V}{m}$$

where $C_0$ and $C_e$ are the UA concentration (μM) at the initial solution and equilibrium adsorption, respectively, V is volume (mL) of the solution, and m is the mass (mg) of adsorbents.

**Adsorption kinetics study**

The mixtures of 3.2 mg δ-$MnO_2$ and 40 mL of 500 μM UA SIF solutions in 50 mL plastic centrifuge tubes were shaken at 180 rpm in a 37 °C constant temperature shaker for 5, 10, 20, 30, 60, 90, 120, 180, 240, and 360 min. The suspension was filtered through a 0.22 μm filter and the concentration ($C_t$) of UA in the filtrate was measured with a microplate reader at 291 nm.

The adsorption capacity ($Q_t$, mg $g^{-1}$) at different incubation times was calculated using the following equation:

$$Q_t = 0.168 * (C_0 - C_t) * \frac{V}{m}$$

where $C_0$ and $C_t$ are the UA concentration (μM) at the initial solution and different incubation times, respectively, V is volume (mL) of the solution, and m is the mass (mg) of δ-$MnO_2$.

The adsorption experimental data was analyzed using pseudo-first-order and pseudo-second-order kinetic models to calculate the kinetics parameters for the removal of UA with δ-$MnO_2$.

The linear equation of pseudo-first-order is expressed as follows:

$$\ln(Q_e - Q_t) = lnQ_e - k_1 t$$

where $Q_e$ and $Q_t$ are the adsorption capacity of UA with δ-$MnO_2$ (mg $g^{-1}$) at equilibrium point and at different incubation time t (min), respectively, and $k_1$ is the adsorption rate constant ($min^{-1}$) for the pseudo-first-order kinetic model.

The pseudo-second-order kinetic rate constant was calculated using the following linear equation:

$$\frac{t}{Q_t} = \frac{1}{k_2 Q_e^2} + \frac{t}{Q_e}$$

where $k_2$ is the adsorption rate constant (g $mg^{-1}$ $min^{-1}$) of the adsorption pseudo-second-order kinetic model.

**Adsorption isotherm study**

A 50 mL plastic centrifuge tube containing 3.2 mg δ-$MnO_2$ and 40 mL SIF solutions of UA with initial concentrations of 400, 500, 600, 700, 800, and 900 μM were prepared, respectively. These tubes were shaken at 180 rpm in a 37 °C constant temperature shaker for 4 h. The sample was filtered through a 0.22 μm filter once the adsorption achieved equilibrium, and the concentration ($C_e$) of UA in the filtrate was quantified with a microplate reader at 291 nm. The adsorption capacity ($Q_e$, mg $g^{-1}$) was calculated according to the equilibrium concentration of UA.

The adsorption process and isotherm parameters for UA adsorption onto δ-$MnO_2$ were described using Langmuir and Freundlich isotherms in this work.

The linearized version of the Langmuir isotherm model is described as follows:

$$\frac{C_e}{Q_e} = \frac{1}{K_L Q_m} + \frac{C_e}{Q_m}$$

where $C_e$ is the equilibrium concentration (mg $L^{-1}$) of UA, $Q_e$ is the adsorption capacity of UA with δ-$MnO_2$ (mg $g^{-1}$) at equilibrium point, $Q_m$ is the maximum adsorption amount (mg $g^{-1}$) of monolayer adsorbed surface of the adsorbent, and $K_L$ is the Langmuir constant (L $mg^{-1}$).

The separation factor or equilibrium parameter ($R_L$) is an essential Langmuir model parameter that is used to determine whether the adsorption is favorable or unfavorable. The values of $R_L$ indicate the adsorption isotherm shapes, and it can be expressed as the following equation:

$$R_L = \frac{1}{1 + K_L C_0}$$

where $K_L$ (L $mg^{-1}$) and $C_0$ (mg $L^{-1}$) are the Langmuir constant and the initial UA concentration, respectively. The adsorption is considered favorable ($0 < R_L < 1$), unfavorable ($R_L > 1$), linear ($R_L$=1), or irreversible ($R_L$ =0).

The linearized form of the Freundlich isotherm model can be written as follows:

$$lnQ_e = lnK_F + \frac{1}{n} lnC_e$$

where $C_e$ is the equilibrium concentration (mg $L^{-1}$) of UA, $Q_e$ is the saturated adsorption capacity of UA with δ-$MnO_2$ (mg $g^{-1}$), and $K_F$ and 1/n are the Freundlich constants that might reflect the adsorption capacity (L $mg^{-1}$) and the adsorption intensity or the surface heterogeneity, respectively. The adsorption is deemed favorable (0 < 1/n < 1), unfavorable (1/n > 1) and irreversible (1/n = 1).

**Adsorption thermodynamics study**

A 50 ml centrifuge tube includes 3.2 mg δ-$MnO_2$ and 40 mL SIF solutions of UA with different initial concentrations. These centrifuge tubes were shaken in a thermostatic shaker at 24, 28, 32, 36 and 40 °C with constant shaking speed at 180 rpm for 4 h. After the completion of adsorption, the suspension was filtered through a 0.22 μm filter. The concentration ($C_e$) of UA in the filtrate was analyzed with a microplate reader and was used to calculate the adsorption amount ($Q_e$, mg $g^{-1}$) at equilibrium point. Then, the Langmuir constant $K_L$ (L $mg^{-1}$) was calculated according to the linearized equation of the Langmuir isotherm model.

The thermodynamic parameters in the adsorption process such as the changes of Gibbs free energy ($\Delta G^0$), the changes of enthalpy ($\Delta H^0$), and the changes of entropy ($\Delta S^0$) were calculated using the Van't Hoff equation.

$$\Delta G^0 = -RTlnK_0$$

$$\Delta G^0 = \Delta H^0 - T\Delta S^0$$

$K_0$ may be obtained by combining above-mentioned Van't Hoff equations, which is given as the following equation:

$$lnK_0 = -\frac{\Delta H^0}{RT} + \frac{\Delta S^0}{R}$$

where R is the gas constant (8.314 J $K^{-1}$ $mol^{-1}$), T is the absolute temperature (Kelvin), and $K_0$ is the thermodynamic equilibrium constant (dimensionless).

The value of $K_0$ was calculated using the following equation:

$$K_0 = \frac{1}{\gamma} * 1000 * K_L * M_w * [Adsorbate]^0$$

where γ is the activity coefficient (dimensionless), $M_w$ is the molecular weight of UA (168.11 g $mol^{-1}$), $[Adsorbate]^0$ is the standard concentration of UA (1 mol $L^{-1}$), $K_L$ is the value of Langmuir equilibrium constant (L $mg^{-1}$) which is the best fitted isotherm model. For diluted solutions, the value of the activity coefficient is unitary, and the activity coefficient is assumed to be 1.

**Adsorption selectivity and interference components**

The mixture of 500 μM UA solutions with 500 μM SIF solutions of different small-molecule organic interfering substances (citric acid, ascorbic acid, adenine, cytosine, uracil, thymine, L-phenylalanine, L-tyrosine, L-cysteine), or 5/20 mM SIF solutions of different flavor or energetic interfering substances (L-glutamate, glucose, sodium oleate, sodium stearate), or 5/50 mM aqueous solutions of different inorganic interfering substances ($NaHCO_3$, $CaCl_2$, $MgCl_2$, NaCl, KCl, $ZnSO_4$, $KH_2PO_4$) were prepared. The centrifuge tubes including 3.2 or 4 mg δ-$MnO_2$ and 40 mL UA solutions with interfering substances were shaken at 180 rpm in a thermostatic shaker at 37 °C for 4 h. The suspension was filtered through a 0.22 μm filter and the equilibrium concentration ($C_e$) of UA, or adenine, cytosine, uracil, or thymine in the filtrate was measured with a microplate reader at 291 nm, or 260, 267, 262, or 264 nm, respectively. The standard curve was plotted using the solution containing interfering substance.

The adsorption capacity (Q, mg $g^{-1}$) of UA onto δ-$MnO_2$ in the interfering substance solutions was calculated using the following equation:

$$Q = 0.168 * (C_0 - C_e) * \frac{V}{m}$$

where $C_0$ and $C_e$ are the UA concentration (μM) at the initial solution and after incubation for 4 h, respectively, V is volume (mL) of the solution, and m is the mass (mg) of δ-$MnO_2$.

The selectivity coefficient (k)[5] of δ-$MnO_2$ for UA to adenine, cytosine, uracil, or thymine can be obtained from the following equation:

$$k = \frac{K_d(UA)}{K_d(\text{adenine, cytosine, uracil, or thymine})}$$

where $K_d$ is the distribution adsorption coefficient, which can be obtained from the following equation:

$$K_d = \frac{C_0 - C_e}{C_e} * \frac{V}{m}$$

where $C_0$ and $C_e$ is the initial and equilibrium concentration (μM) of adenine, cytosine, uracil, thymine or UA, respectively, V is the solution volume (mL), and m is the mass (mg) of δ-$MnO_2$.

**Desorption of UA in δ-$MnO_2$**

A 250 mL bottle with blue cap including 40 mg δ-$MnO_2$ and 200 mL UA SIF solution with an initial concentration of 500 μM was shaken at 180 rpm in a 37 °C shaker for 4 h. The black solid was collected from the mixture by centrifugation. The solid was put in a 0.22 μm filter equipped with a 1 mL syringe and washed 8 times with a 1 mL 0.1 M sodium hydroxide solution. The UA concentration in the filtrate was determined with a microplate reader at 291 nm.

**UA oxidation by δ-$MnO_2$**

The UA was dissolved in a dilute sodium hydroxide solution and then the pH was adjusted to 7 using dilute hydrochloric acid solution. The mixture of δ-$MnO_2$ and UA aqueous solution was shaken at 180 rpm in a 37 °C shaker for 4 h. The mixture was filtered through a 0.22 μm filter and the chemicals of UA degradation catalyzed by δ-$MnO_2$ in the filtrate were investigated using a mass spectrometer (LTQ-XL, Thermo Fisher Scientific).

**Manganese release of δ-$MnO_2$ in simulated GI fluid**

A 10 mL centrifuge tube with 2 g $L^{-1}$ δ-$MnO_2$ in SGF or SIF solution was shaken at 180 rpm in a 37 °C shaker for 4 h. The mixture was filtered through a 0.22 μm filter and the manganese content in filtrate was determined by inductively coupled plasma optical emission spectroscopy (ICP-OES) on a Thermo Scientific iCAP 7600 spectrometer.

**Animals**

All animal studies were carried out in accordance with the guidelines of the Institutional Animal Care and Use Committee of Shanghai Jiao Tong University. In this work, C57BL/6J or KM mice (6 weeks) were obtained from SPF (Beijing) Biotechnology Co., Ltd. (Beijing, China) for *in vivo* toxicity, magnetic resonance imaging, or drug-induced hyperuricemia model studies. A total of 24 male C57BL/6J Uox KO mice (8 to 10 weeks) generated at Changzhou Cavens Lab Aminal Co., Ltd. (Changzhou, China) were used as hyperuricemia mouse model. The mice were maintained on a 12-hour light/12-hour dark cycle and acclimatized for one week prior to the experiment.

***In vivo* toxicity and manganese biodistribution study**

To evaluate the acute toxicity of δ-$MnO_2$, C57BL/6J mice were gavaged with 0.2 mL of δ-$MnO_2$ aqueous suspension at a dosage of 0, 0.25, 0.5, 1, and 2 g $kg^{-1}$ for 14 days. Body weight change, water and food consumption, mortality, behavioral manifestations such as irritability and agitation were all meticulously tracked. On day 15, whole blood samples and serum were collected from some mice for routine blood test, biochemical indexes and blood cell manganese content analysis. The mice were euthanized and dissected after blood collection. Heart, liver, spleen, lung, kidney, brain, stomach, 1-2 cm intestine, 1-2 cm colon, and hair were collected for manganese content measurement or hematoxylin and eosin staining. From the $15^{th}$ day to the $21^{st}$ day, the remaining mice were fed without any treatment to test the recovery of liver manganese content after 14 days repeated administration of δ-$MnO_2$. The manganese content in biological tissues was investigated using an inductively coupled plasma mass spectrometry (ICP-MS) on a PerkinElmer NexION2000-Flexar20 HPLC spectrometer.

**Magnetic resonance imaging (MRI) of GI tract with δ-$MnO_2$ gavage**

Mn is a paramagnetic MRI contrast agent in δ-$MnO_2$ that allows real-time imaging of the δ-$MnO_2$ excretion process through the gastrointestinal tract. The health C57BL/6J mice were orally administered δ-$MnO_2$ (250 mg $kg^{-1}$) aqueous solution. After sedation with isoflurane, T1 and T2 weighted images of mice at 1 h were acquired using a magnetic resonance imaging instrument (Bruker-Biospec 94/30 USR).

***In vivo* UA-lowering experiment**

There are three mice models of hyperuricemia used to investigate the in vivo UA-lowering effects of δ-$MnO_2$. For the first model, male KM mice were used to construct the hyperuricemia induced by intraperitoneal injection of potassium oxonate (300 mg $kg^{-1}$) and intragastric administration of UA (30 mg $kg^{-1}$). Allopurinol (6 or 60 mg $kg^{-1}$), or δ-$MnO_2$ at various doses (30, 50, 125, 250 mg $kg^{-1}$) were orally administered. The blood sample was collected from the submandibular vein into a micro blood collection tube at 50 or 60 min after intraperitoneal injection and centrifuged at 3,000 g for 15 min at 4 °C. The serum UA was measured using UA content assay kit (Solarbio, Beijing, China).

For the second model, male KM mice were used to construct the hyperuricemia induced by potassium oxonate (4.4%, w/w) and adenine (0.44%, w/w) in basal diet for 14 days. The basal diet contained 57.3% (w/w) chow powder rodent diet, 11.4% (w/w) peanut butter, 11.4% (w/w) pig intestine protein, 11.4% (w/w) sucrose, and 8.5% (w/w) olive oil and was provided *ad libitum*. The positive drug was administered through drinking water with 50 mg $L^{-1}$ allopurinol. δ-$MnO_2$ (1.16%, w/w) were administered with the basal diet containing hyperuricemia-inducing drugs. Mice were put in metabolic cages at 13 d of the treatment period to collect 24-hour urine for the analysis of UA and creatinine levels. 1-2 drops of an 8 M NaOH aqueous solution were added to the urine collection containers to prevent precipitation of UA salts. The blood sample was collected from the submandibular vein into a micro blood collection tube at 14 d of the treatment period and centrifuged at 3,000 g for 15 min at 4 °C for the analysis of UA levels and creatinine concentration. The UA and creatinine levels were measured using UA content assay kit (Solarbio, Beijing, China), and creatinine assay kit (Shanghai Yuanye Bio-Technology Co., Ltd, Shanghai, China), respectively.

For the third model, the 24 male C57BL/6J Uox KO mice were randomly divided into three groups: control (n = 8); 40 mg $kg^{-1}$ $d^{-1}$ allopurinol positive group (n = 8); 250 mg $kg^{-1}$ $d^{-1}$ δ-$MnO_2$ treatment group (n = 8). During the experimental period, all mice were gavaged twice a day and their body weights were monitored every day. Mice were put in metabolic cages at 1 week after gavage during the treatment period to collect the 24-hour urine for the analysis of urinary UA and creatinine. 1-2 drops of an 8 M NaOH aqueous solution were added to the urine collection containers to prevent precipitation of UA salts. Following the collection of urine, 200 μL blood sample was collected from the submandibular vein into a micro blood collection tube and centrifuged at 3,000 g for 15 min at 4 °C for the analysis of serum UA and creatinine. The UA and creatinine levels were measured using above-mentioned kits.

**Calculation of fractional excretion of uric acid (FEUA) and uric acid clearance rate (CUA)**

The fractional excretion of uric acid (FEUA) is defined as the percentage of UA filtered through the glomeruli that is excreted in the urine and is calculated using the following equation:

$$FEUA\ (\%) = \frac{UUA \times SCRE}{SUA \times UCRE} \times 100$$

where UUA and SUA are the urine and serum UA (mg $dL^{-1}$), respectively; SCRE and UCRE are the serum and urine creatinine (mg $dL^{-1}$), respectively.

The uric acid clearance rate (CUA) was calculated from the following equation:

$$CUA\ \left(\frac{mL}{h}\right) = \frac{UUA \times UV}{24 \times SUA}$$

where UUA, UV, and SUA are the 24-h urinary levels of UA (μM), urinary volume (mL), and serum UA (μM), respectively.

### Supporting Figures

**a**

This study primarily aims to understand the **adsorption** mechanism of **urea**, **creatinine**, and **uric acid** on spherical **activated carbon**. The adsorption of urea, creatinine, and uric acid onto spherical activated carbon underwent with a pseudo-second-order rate and in accordance with the Halsey formula. The adsorption of urea may have occurred because of interaction between the urea dipole and the dipole induced in the porous surface by the urea as well as because of dipole–dipole interaction between surface oxygen groups on the spherical activated carbon surface and the urea. The interaction among urea molecules, such as hydrogen bonding, induced multilayer adsorption. The adsorption model of creatinine was similar to that of urea. Because uric acid molecules are very strongly hydrophobic, their adsorption onto spherical activated carbon is caused by attractive forces between the hydrophobic surface of activated carbon and the similarly hydrophobic uric acid molecules, in addition to van der Waals forces. Moreover, uric acid molecules adsorbed onto spherical activated carbon and uric acid molecules in water are considered to undergo additional multilayer adsorption because of hydrophobic interactions. (Kameda, T., Horikoshi, K., Kumagai, S., Saito, Y. & Yoshioka, T. Adsorption of urea, creatinine, and uric acid onto spherical activated carbon. *Sep. Purif. Technol.* **237**, 116367 (2020).)

**b**

Development of novel nanomaterials for biosensors has intrigued widespread interest. Here, we report a method to graft the redox-active dye Methylene Blue (MB) onto **molybdenum disulfide ($MoS_2$)** nanosheet surface via electrostatic and π-stacking interaction. The **adsorption** of MB on nanosheets was investigated by atomic force microscopy (AFM), which proved that the adsorption isotherm fits a Temkin not a Langmuir model. After studying the electrochemical properties of MB-decorated $MoS_2$ nanocomposite ($MoS_2$@MB) on a glassy carbon electrode (GCE), an electrochemical sensor for microRNA-21 detection was designed. The modified GCE can quantify microRNA-21 in concentrations as low as 68 fM, typically at a working potential of −0.28 V (vs. SCE). The same modified electrode also shows outstanding electrocatalytic ability towards individual and simultaneous determination of dopamine (DA) and **uric acid (UA)** with electrochemical peaks at 0.16 V (DA) and 0.45 V (UA). The detection limits for simultaneous determination are 0.58 μM for DA and 0.91 μM for UA, respectively. (Su, S. et al. A molybdenum disulfide@Methylene Blue nanohybrid for electrochemical determination of microRNA-21, dopamine and uric acid. *Microchim. Acta* **186**, 607 (2019).)

**c**

This study sought to evaluate the role of N-acetyl cysteine (NAC) on ibuprofen-induced hepatotoxicity in rats. The rats were divided into six groups. Group 1 (control group) received carboxy-**methyl cellulose**. Group 2 (untreated group) was given ibuprofen. Group 3 administrated with ibuprofen and silymarin. Groups 4, 5, and 6 were given ibuprofen and NAC. We assessed histopathological examinations and serum biochemical parameters such as alkaline phosphatase (ALP), glutamate pyruvate transaminase (GPT), urea, glutamate oxaloacetate transaminase (GOT), uric acid, lipid profile, serum interleukin-1 beta (IL-1β), catalase (CAT), superoxide dismutase (SOD), vitamin C (Vit C), protein carbonyl (PC), and ferric reducing antioxidant power (FRAP). Group 2 revealed a remarkable elevation ($p < 0.05$) in serum ALP, GPT, GOT, lipid profile (except HDL-C), PC, **uric acid**, MDA, serum IL-1β, and its hepatic gene expressions relate to group 1. Also in group 2, plasma FRAP and liver CAT, SOD, and Vit C significantly reduced ($p < 0.05$) as opposed to group 1. Nevertheless, NAC and silymarin led to improvement in the above parameters in contrast with those of group 2 in the treated rats. Our results confirmed protective effects of NAC on ibuprofen-induced hepatoxicity in male rats through increasing parameters such as CAT, SOD, Vit C, and FRAP. (Satvati, M., Salehi-Vanani, N., Nouri, A. & Heidarian, E. Protective effects of N-acetyl cysteine against oxidative stress in ibuprofen-induced hepatotoxicity in rats. *Comp. Clin. Pathol.* **31**, 293–301 (2022).)

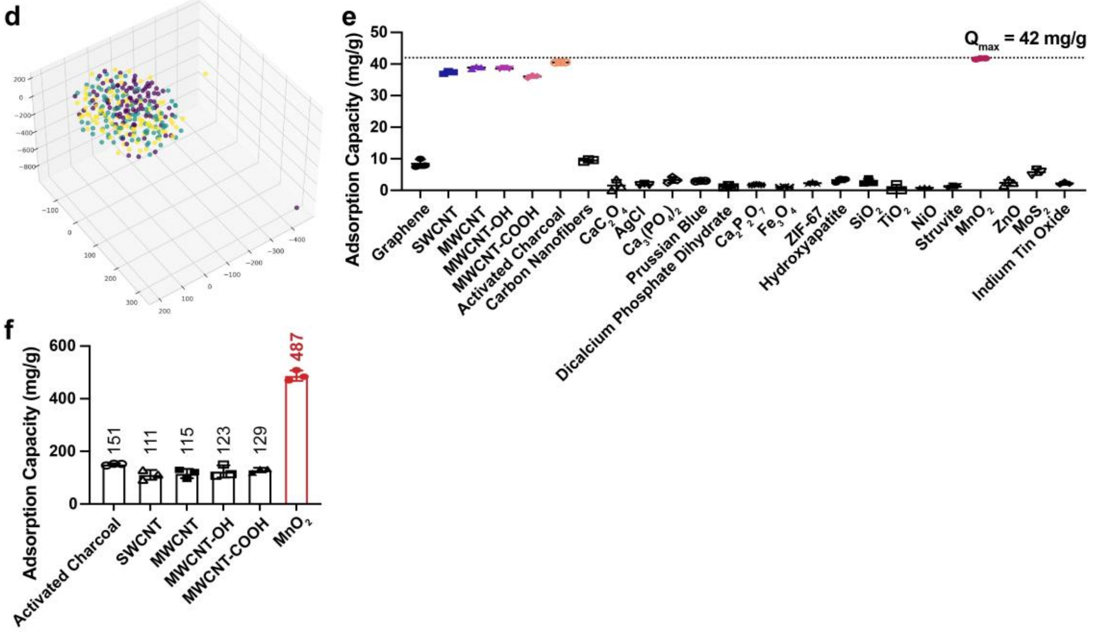


**Figure S1. NLP and experimental screening for UA adsorption material discovery. a**,**b**, The examples of literature abstracts with irrelevant co-occurring entities (**a**, blue words) and distant context (**b**) for the semantics of uric acid, adsorption, and material entities. **c**, The example of literature abstract including material entity (green words) with low cosine similarity.

**d**, t-SNE plots of 288 chemical entities with higher cosine similarity from UA related literatures. **e**, Adsorption capacities of UA onto 24 types of materials at the dosage of 2 g $L^{-1}$ in SIF. **f**, Adsorption capacities of UA onto activated charcoal, single walled carbon nanotubes (SWCNT), multi-walled carbon nanotubes (MWCNT), hydroxylated multi-walled carbon nanotubes (MWCNT-OH), carboxylated multi-walled carbon nanotubes (MWCNT-COOH), and $MnO_2$ at the dosage of 0.1 g $L^{-1}$ in SIF.

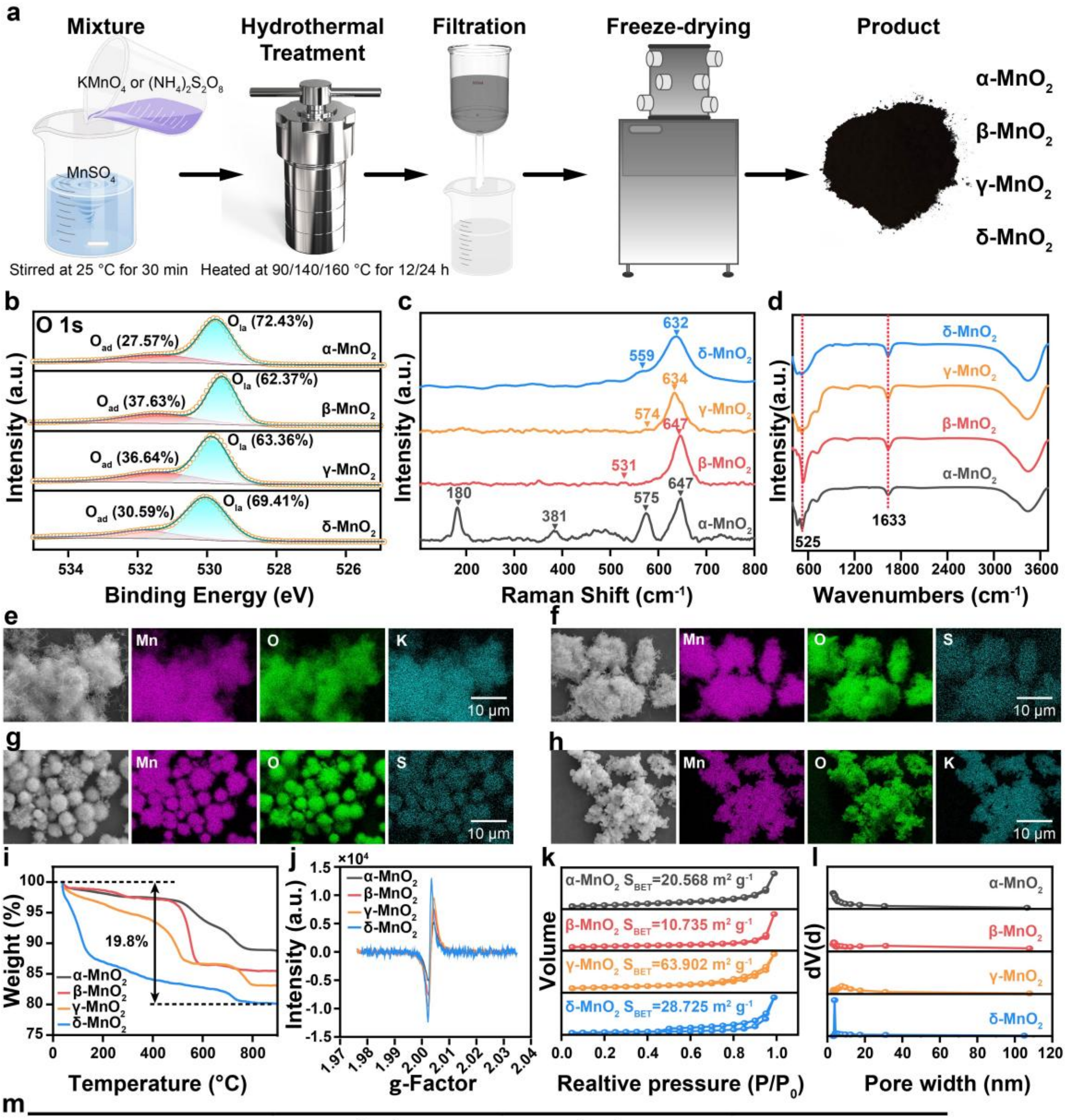


m

| Materials | SSA ($m^2$ $g^{-1}$) | $V_{tot}$ ($m^3$ $g^{-1}$) | Pore size (nm) |
|---|---|---|---|
| **α-$MnO_2$** | 20.6 | 0.08 | 3.06 |
| **β-$MnO_2$** | 10.7 | 0.06 | 3.41 |
| **γ-$MnO_2$** | 63.9 | 0.29 | 3.06 |
| **δ-$MnO_2$** | 28.7 | 0.18 | 30.06 |

**Figure S2. Synthesis and characterizations of α, β, γ, δ-$MnO_2$. a**, The preparation procedure of $MnO_2$ by hydrothermal approach at different conditions. **b**, The XPS spectra of O 1s for the as-synthesized $MnO_2$. Peaks with binding energies at 529.6-529.8 and 531.4-531.8 eV in the XPS spectra of O 1s can be attributed to lattice oxygen ($O_{la}$) and surface adsorption oxygen ($O_{ad}$), respectively. **c**, The Raman spectra for the as-synthesized α, β, γ, δ-$MnO_2$. α-$MnO_2$ featured four major peaks located at approximately 180 $cm^{-1}$, 381 $cm^{-1}$, 575 $cm^{-1}$ and 647 $cm^{-1}$. The peak at 575 $cm^{-1}$ was definitely assigned to the deformation modes of the Mn-O-Mn chain in the $MnO_2$ octahedral lattice while the peak at 647 $cm^{-1}$ was assigned to the Mn-O stretching modes. The peak at 180 $cm^{-1}$ of α-$MnO_2$ represented an external mode that is derived from the translational motion of basic [$MnO_6$] octahedral units. β-$MnO_2$ demonstrated two main peaks at around 531 $cm^{-1}$ and 647 $cm^{-1}$. The peak at 531 $cm^{-1}$ corresponded to the Mn-O stretching of the [$MnO_6$] octahedral unit and the peak at 647 $cm^{-1}$ was indexed to the characteristic $A_{1g}$ mode. The γ-$MnO_2$ also displayed two major peaks situated at about 574 $cm^{-1}$ and 634 $cm^{-1}$. The former peak at 574 $cm^{-1}$ suggested a well-developed orthorhombic structure with a (2×1) tunnel in γ-$MnO_2$ and the latter peak at 634 $cm^{-1}$ was ascribed to the stretching mode of the Mn-O bond in the [$MnO_6$] octahedral unit. The Raman spectrum of δ-$MnO_2$ with two bands appear at ~632 $cm^{-1}$ and ~559 $cm^{-1}$, respectively. The former arises from the symmetric stretching vibrations of Mn-O bonds in $MnO_6$ groups, while the latter is attributed to Mn-O stretching in the basal plane of $MnO_6$ sheet. **d**, The FTIR spectra of the as-synthesized α, β, γ, δ-$MnO_2$. The broad peak at 525 $cm^{-1}$ can be assigned to Mn-O vibration, and a sharp peak at 1633 $cm^{-1}$ can be assigned to Mn-O-Mn vibration. **e-h**, Energy dispersive spectroscopy (EDS) analysis of α-$MnO_2$ (**e**), β-$MnO_2$ (**f**), γ-$MnO_2$ (**g**), and δ-$MnO_2$ (**h**). Scale bar: 10 μm. Mn, O, K, or S elements are uniformly distributed within the particles. **i**, The thermograms of thermogravimetric analysis for $MnO_2$. **j**, EPR spectra at 100 K of oxygen vacancy defects for $MnO_2$. **k-m**, $N_2$ adsorption and desorption isotherms (**k**), pore size distributions (**l**), and the values of specific surface area (SSA/BET), total pore volume ($V_{tot}$) and pore size (**m**) for the as-synthesized α, β, γ, δ-$MnO_2$. The shapes of the isotherms are type IV for the property of mesoporous materials.

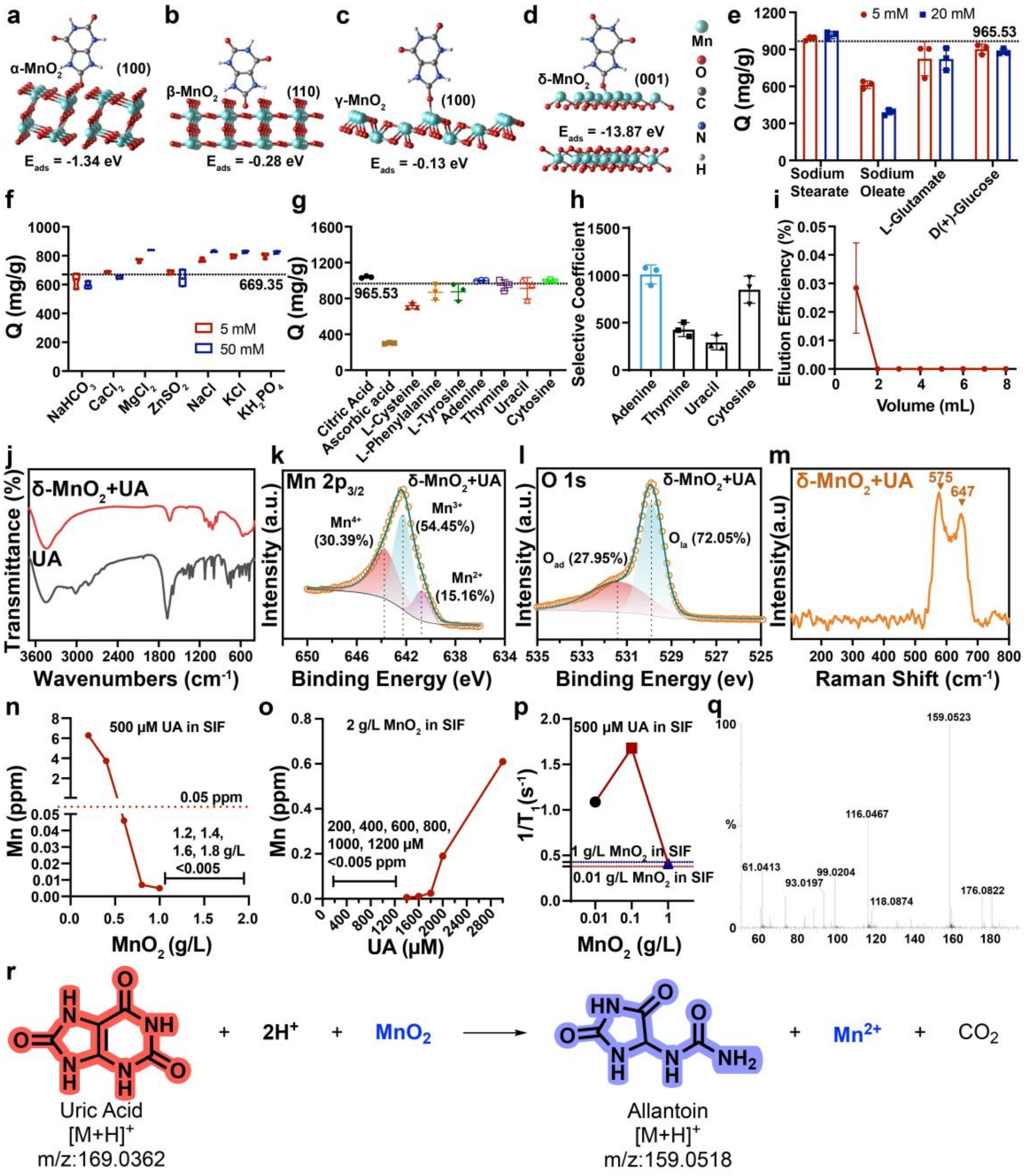


**Figure S3. The adsorption and oxidation studies of UA and $MnO_2$. a-d**, The UA adsorption energies of α, β, γ, δ-$MnO_2$ using DFT calculation. DFT calculation was employed in the Cambridge Sequential Total Energy Package (CASTEP) code. The exchange-correction function was described by the generalized gradient approximation (GGA) within the Perdew-Burke-Ernzerhof (PBE) function with Hubbard U corrections (PBE + U). The Hubbard U value is set to 1.6 eV to correct the 3d orbital electronic structure of Mn atoms. The energy cutoff of the plane waves was set to 450 eV. The Brillouin zone sampled with 1 × 1 × 1, 1 × 1 × 1, 1 × 1 × 1 and 2 × 2 × 1 k-points mesh by the Monkhorst-Pack method were used for α-, β-, γ- and δ-$MnO_2$, respectively. For the geometry optimization, convergence thresholds were set to 5.0 × $10^{-4}$ Ha for energy, 0.001 Ha/Å for force, and 0.005 Å for displacement. To eliminate artificial interactions between adjacent images, a vacuum space of ~15 Å was adopted. The UA structure was optimized in the same unit cell as the $MnO_2$ geometric structure optimization, and the k-point was placed at the Gamma point, while the other parameters were consistent with the $MnO_2$ optimization parameters. The adsorption energies ($E_{ads}$) of UA adsorbed on the α-, β-, γ- and δ-$MnO_2$ monolayers were calculated as follows: $E_{ads} = E_{total} - E_{UA} - E_{adsorbent}$, where $E_{adsorbent}$, $E_{UA}$, and $E_{total}$ refer to the energies of the α-, β-, γ- and δ-$MnO_2$ substrates, UA, and α-, β-, γ- and δ-$MnO_2$ substrates with adsorbed UA, respectively. **e**, The adsorption capacities of UA onto δ-$MnO_2$ in SIF (pH 6.8) with 5 or 20 mM interfering substances including sodium stearate, sodium oleate, L-glutamate, or D-(+)-glucose (n = 3 replicates). The value of dotted line represents the UA adsorption capacity in SIF (pH 6.8). **f**, The relationship of the UA adsorption capacity onto δ-$MnO_2$ and the inorganic salts including $NaHCO_3$, $CaCl_2$, $MgCl_2$, $ZnSO_4$, NaCl, KCl, or $KH_2PO_4$ at concentrations of 5 or 50 mM (pH 6.8, n = 3 replicates). The value of dotted line represents the UA adsorption capacity in water (pH 6.8, adjusted with NaOH). **g**, The adsorption capacities of UA onto δ-$MnO_2$ in SIF (pH 6.8) in the presence of interfering substances including citric acid, ascorbic acid, L-cysteine, L-phenylalanine, L-tyrosine, adenine, thymine, uracil, or cytosine with the consistent concentration (500 μM) of UA (n = 3 replicates). The value of dotted line shows the UA adsorption capacity in SIF (pH 6.8). **h**, Selectivity coefficient of δ-$MnO_2$ for UA with respect to adenine, thymine, uracil, and cytosine (n = 3 replicates). The selectivity coefficient is defined as the ratio of the equilibrium constants for the two competing species and is a measure of the relative affinity of δ-$MnO_2$ for UA over adenine, thymine, uracil, or cytosine (the higher value, the stronger relative affinity of δ-$MnO_2$ to UA). **i**, Elution curve of UA using NaOH aqueous solution (0.1 M) as the eluent after UA adsorption onto δ-$MnO_2$ solid. The dynamic desorption curve shows that almost no UA was eluted. **j-m**, FTIR (**j**), Mn $2p_{3/2}$ (**k**) and O 1s (**l**) XPS, and Raman (**m**) spectra of the δ-$MnO_2$ adsorbed UA sample. **n**, The manganese content in the filtrate after UA at 500 μM in SIF adsorption onto δ-$MnO_2$ with different dosages (0.2, 0.4, 0.6, 0.8, 1.0, 1.2, 1.4, 1.6, and 1.8 g/L) for 4 h at 37 °C. **o**, The manganese content in the filtrate after UA adsorption onto 2 g/L δ-$MnO_2$ in SIF at different concentrations (200, 400, 600, 800, 1000, 1200, 1400, 1600, 1800, 2000, and 3000 μM) for 4 h at 37 °C. **p**, T1-MR signals of 500 μM UA and δ-$MnO_2$ with different dosages (0.01, 0.1, and 1 g/L) in SIF. The red and blue dotted lines represent the T1-MR signals of pure δ-$MnO_2$ in SIF at 0.01 and 1 g/L dosages, respectively. The magnetic resonance signal of $1/T_1$ initially increases and subsequently decreases with the increase of δ-$MnO_2$ dosage, owing the destruction of a portion of $MnO_2$ into paramagnetic $Mn^{2+}$ within the appropriate concentration range. **q**, Mass spectrum of allantoin found in the filtrate after UA adsorption onto δ-$MnO_2$. **r**, The proposed reaction equation and molecular information for manganese dioxide reacting with UA in aqueous solution to form allantoin.

**a**

| Source | Sequential p-value | Lack of Fit p-value | Adjusted R² | Predicted R² | |
|---|---|---|---|---|---|
| Linear | < 0.0001 | 0.0434 | 0.9222 | 0.8975 | |
| **2FI** | **0.0175** | **0.1062** | **0.9511** | **0.9142** | **Suggested** |
| Quadratic | 0.1947 | 0.1296 | 0.9577 | 0.8880 | |
| Cubic | 0.1147 | 0.2664 | 0.9770 | 0.6671 | **Aliased** |

**b**

| | | | |
|---|---|---|---|
| **Std. Dev.** | 43.10 | **R²** | 0.9680 |
| **Mean** | 552.18 | **Adjusted R²** | 0.9511 |
| **C.V. %** | 7.80 | **Predicted R²** | 0.9142 |
| | | **Adeq Precision** | 30.3680 |

**c**

| Source | Sum of Squares | df | Mean Square | F-value | p-value | |
|---|---|---|---|---|---|---|
| **Model** | 1.067E+06 | 10 | 1.067E+05 | 57.43 | < 0.0001 | significant |
| A-Uric acid | 76729.15 | 1 | 76729.15 | 41.31 | < 0.0001 | |
| B-pH | 8.493E+05 | 1 | 8.493E+05 | 457.27 | < 0.0001 | |
| C-Adsorbent | 27649.96 | 1 | 27649.96 | 14.89 | 0.0011 | |
| D-Time | 74464.05 | 1 | 74464.05 | 40.09 | < 0.0001 | |
| AB | 26073.68 | 1 | 26073.68 | 14.04 | 0.0014 | |
| AC | 41.76 | 1 | 41.76 | 0.0225 | 0.8824 | |
| AD | 1463.54 | 1 | 1463.54 | 0.7879 | 0.3858 | |
| BC | 8460.14 | 1 | 8460.14 | 4.55 | 0.0461 | |
| BD | 2499.06 | 1 | 2499.06 | 1.35 | 0.2604 | |
| CD | 54.41 | 1 | 54.41 | 0.0293 | 0.8659 | |
| **Residual** | 35290.92 | 19 | 1857.42 | | | |
| Lack of Fit | 31689.72 | 14 | 2263.55 | 3.14 | 0.1062 | not significant |
| Pure Error | 3601.20 | 5 | 720.24 | | | |
| **Cor Total** | 1.102E+06 | 29 | | | | |

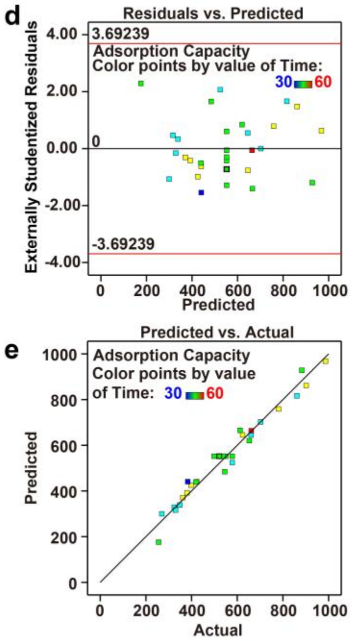


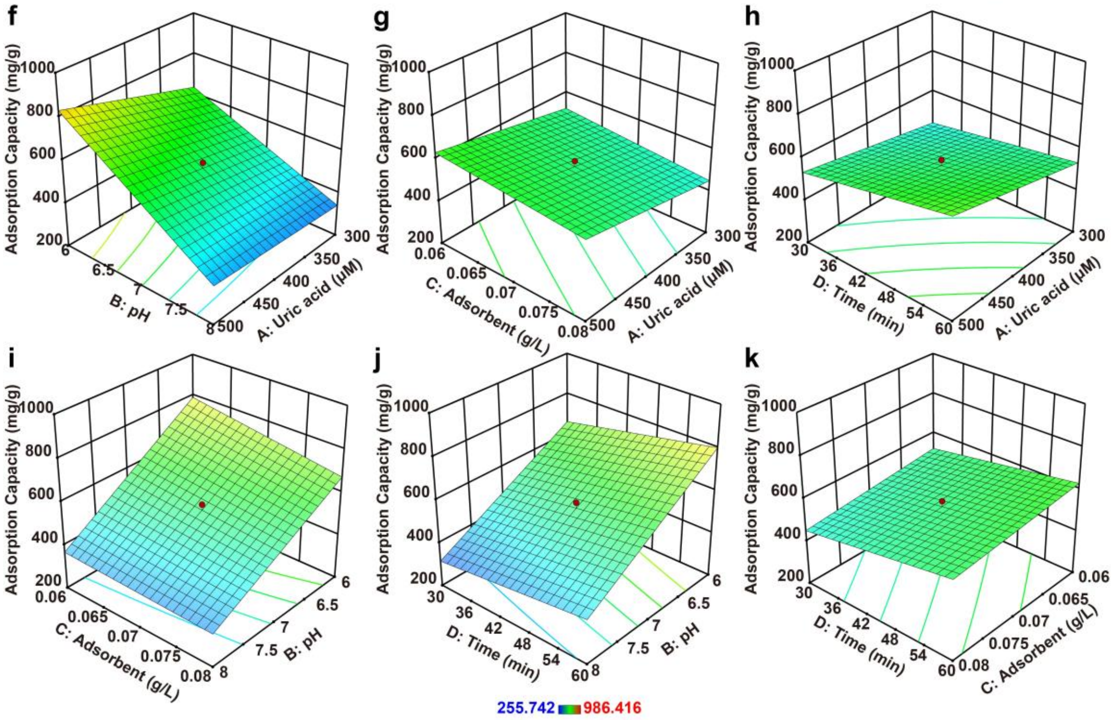


**Figure S4. Optimization of UA adsorption conditions by RSM-CCD. a**, Model statistic results of fit summary in RSM-CCD design for UA removal. Two factor interaction (2FI) model was significant for the adsorption of UA on δ-$MnO_2$. **b**, The results of fit statistics in RSM-CCD design. The accuracy of the 2FI regression model is evaluated by regression coefficient ($R^2$, 0.9680), adjusted $R^2$ (0.9511), and predicted $R^2$ (0.9142). The difference of adjusted and predicted $R^2$ is less than 0.2, indicating that the fit between the experimental and predicted data is desirable. The value of Adeq Precision for the model is 30.37 (adeq>4), which indicates that the signal to noise ratio of the model is in an ideal range. **c**, Results of ANOVA analysis for 2FI model of UA adsorption onto δ-$MnO_2$. The F value of the 2FI model is 57.43, indicating that the model item is significant and there is only a 0.01% chance that could occur due to noise. The lack of fit F value for 2FI model is 3.14 and the probability value (P value) higher than 0.05 were not significant, demonstrating the validity of the model. Model terms with P value less than 0.05, including A (initial UA concentration), B (pH), C (adsorbent dose) and D (contact time), AB, and BC significantly affected the response Y (UA adsorption capacity). The final RSM model (in coded form) for UA adsorption is $\mathrm{Y} = 552.18 + 56.54 * \mathrm{A} - 188.12 * \mathrm{B} - 33.94 * \mathrm{C} + 55.70 * \mathrm{D} - 40.37 * \mathrm{AB} + 1.62 * \mathrm{AC} + 9.56 * \mathrm{AD} + 22.99 * \mathrm{BC} - 12.50 * \mathrm{BD} + 1.84 * \mathrm{CD}$. In the equation, terms with positive coefficient signs infer positive effects on adsorption, whereas terms with negative signs reveal negative effects on adsorption. These results demonstrated that pH, adsorbent dose, the interactions between pH and initial UA concentration or contact time negatively affected the adsorption capacity of UA, whereas initial UA concentration, contact time, the interactions between initial UA concentration and adsorbent dose or contact time, and the interactions between adsorbent dose and pH or contact time showed positive effects. **d**, Plots of the residuals and predicted adsorption capacity values in RSM-CCD design for UA removal. **e**, Plots of the predicted and the actual experimental adsorption capacity values for UA adsorption onto δ-$MnO_2$. There were no outliers in the plots of externally studentized residuals against predicted adsorption capacity. The maximum deviation of the predicted and actual values of UA adsorption capacity is less than 4%, revealing a good correlation between the mathematically calculated values and the experimental values. **f-k**, Three-dimensional surface and contour plots in RSM-CCD design of the interactions among different process variables for UA adsorption capacity onto δ-$MnO_2$: (**f**) pH and initial UA concentration, (**g**) adsorbent dose and initial UA concentration, (**h**) contact time and initial UA concentration, (**i**) adsorbent dose and pH, (**j**) contact time and pH, (**k**) contact time and adsorbent dose. The response is displayed in accordance with the two variable parameters when the other two parameters are constant.

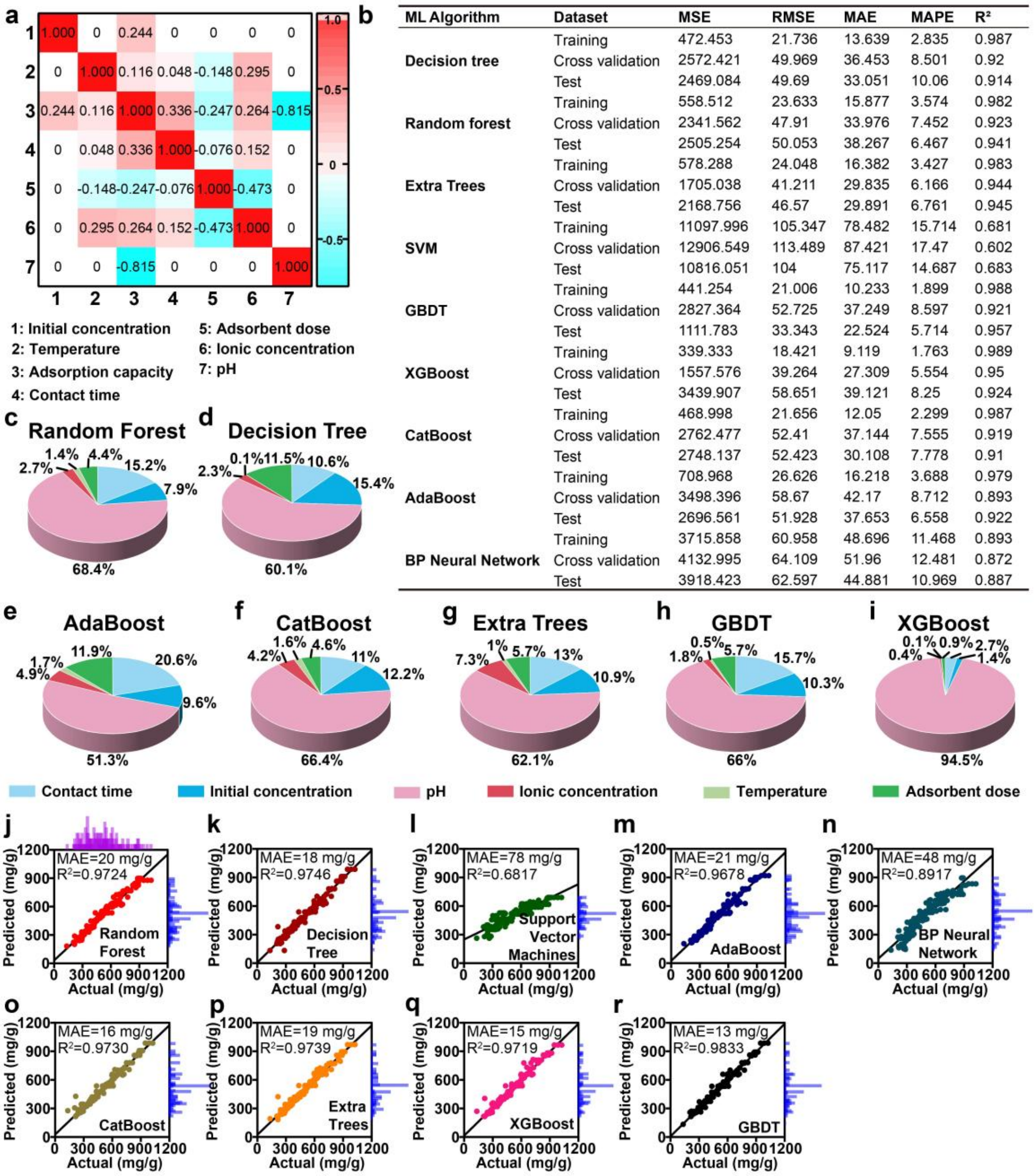


| ML Algorithm | Dataset | MSE | RMSE | MAE | MAPE | R² |
|---|---|---|---|---|---|---|
| Decision tree | Training | 472.453 | 21.736 | 13.639 | 2.835 | 0.987 |
| | Cross validation | 2572.421 | 49.969 | 36.453 | 8.501 | 0.92 |
| | Test | 2469.084 | 49.69 | 33.051 | 10.06 | 0.914 |
| Random forest | Training | 558.512 | 23.633 | 15.877 | 3.574 | 0.982 |
| | Cross validation | 2341.562 | 47.91 | 33.976 | 7.452 | 0.923 |
| | Test | 2505.254 | 50.053 | 38.267 | 6.467 | 0.941 |
| Extra Trees | Training | 578.288 | 24.048 | 16.382 | 3.427 | 0.983 |
| | Cross validation | 1705.038 | 41.211 | 29.835 | 6.166 | 0.944 |
| | Test | 2168.756 | 46.57 | 29.891 | 6.761 | 0.945 |
| SVM | Training | 11097.996 | 105.347 | 78.482 | 15.714 | 0.681 |
| | Cross validation | 12906.549 | 113.489 | 87.421 | 17.47 | 0.602 |
| | Test | 10816.051 | 104 | 75.117 | 14.687 | 0.683 |
| GBDT | Training | 441.254 | 21.006 | 10.233 | 1.899 | 0.988 |
| | Cross validation | 2827.364 | 52.725 | 37.249 | 8.597 | 0.921 |
| | Test | 1111.783 | 33.343 | 22.524 | 5.714 | 0.957 |
| XGBoost | Training | 339.333 | 18.421 | 9.119 | 1.763 | 0.989 |
| | Cross validation | 1557.576 | 39.264 | 27.309 | 5.554 | 0.95 |
| | Test | 3439.907 | 58.651 | 39.121 | 8.25 | 0.924 |
| CatBoost | Training | 468.998 | 21.656 | 12.05 | 2.299 | 0.987 |
| | Cross validation | 2762.477 | 52.41 | 37.144 | 7.555 | 0.919 |
| | Test | 2748.137 | 52.423 | 30.108 | 7.778 | 0.91 |
| AdaBoost | Training | 708.968 | 26.626 | 16.218 | 3.688 | 0.979 |
| | Cross validation | 3498.396 | 58.67 | 42.17 | 8.712 | 0.893 |
| | Test | 2696.561 | 51.928 | 37.653 | 6.558 | 0.922 |
| BP Neural Network | Training | 3715.858 | 60.958 | 48.696 | 11.468 | 0.893 |
| | Cross validation | 4132.995 | 64.109 | 51.96 | 12.481 | 0.872 |
| | Test | 3918.423 | 62.597 | 44.881 | 10.969 | 0.887 |

**Figure S5. Machine learning for the adsorption of UA onto δ-$MnO_2$. a**, PCC matrix for the full set of features of the study of UA adsorption onto δ-$MnO_2$. 1: initial UA concentration (μM); 2: temperature (°C); 3: adsorption capacity (mg $g^{-1}$); 4: contact time (min); 5: adsorbent dose (g $L^{-1}$); 6: ionic concentration (mM); 7: pH (dimensionless). **b**, A comparison of the efficiency of the nine machine learning algorithms in predicting the adsorption capacity of UA onto δ-$MnO_2$. It is noteworthy that all ML methods except SVM and BP neural network have good prediction performance. **c-i**, The pie charts depict the relative proportion for feature (contact time, initial concentration, pH, ionic concentration, temperature, and adsorbent dose) importance of UA adsorption onto δ-$MnO_2$ derived from machine learning algorithms of the Random Forest (**c**), Decision Tree (**d**), AdaBoost (**e**), CatBoost (**f**), Extra Trees (**g**), GBDT (**h**)

XGBoost (**i**). All the results indicated that solution pH was the most significant factor for the adsorption capacity variation (proportion > 50%), which was consistent with the findings of RSM. **j-r**, Parity plots between the experimental and predicted adsorption capacities from five-fold cross-validation of Random Forest (**j**), Decision Tree (**k**), Support Vector Machines (**l**), AdaBoost (**m**), BP Neural Network (**n**), CatBoost (**o**), Extra Trees (**p**), XGBoost (**q**) and GBDT (**r**) algorithms. Histograms on the axis illustrate distributions of actual or predicted adsorption capacity values. The GBDT model with the highest $R^2$ of 0.9833 and the lowest MAE of 12.747 mg/g provided the most accurate prediction of the data. These data suggest that the GBDT model is able to predict the behavior of the UA adsorption process under different conditions.

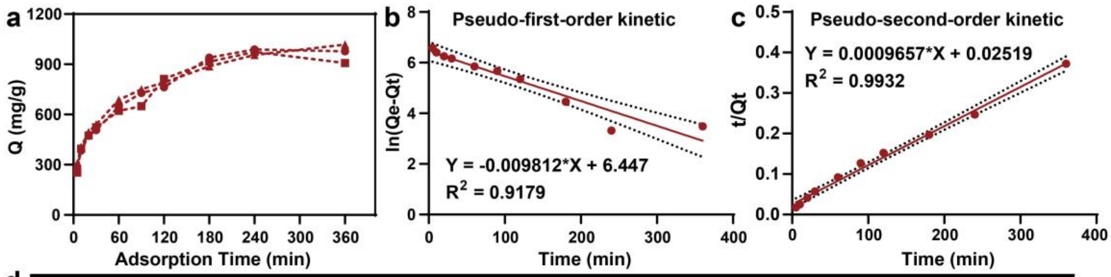


d

| Pseudo-first-order model | | | Pseudo-second-order model | | |
|---|---|---|---|---|---|
| $Q_e$ (mg/g) | $k_1$ | $R^2$ | $Q_e$ (mg/g) | $k_2$ | $R^2$ |
| 630.81 | 0.00981 | 0.9179 | 1035.52 | $3.702\times10^{-5}$ | 0.9932 |

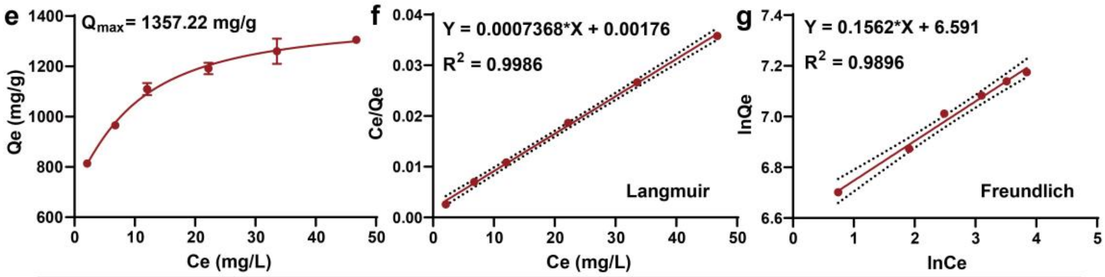


h

| Langmuir model | | | | Freundlich model | | |
|---|---|---|---|---|---|---|
| $Q_m$ (mg/g) | $K_L$(L/mg) | $R^2$ | $R_L$ | n | $K_F$ | $R^2$ |
| 1357.22 | 0.4186 | 0.9986 | 0.0156-0.0343 | 6.402 | 728.51 | 0.9896 |

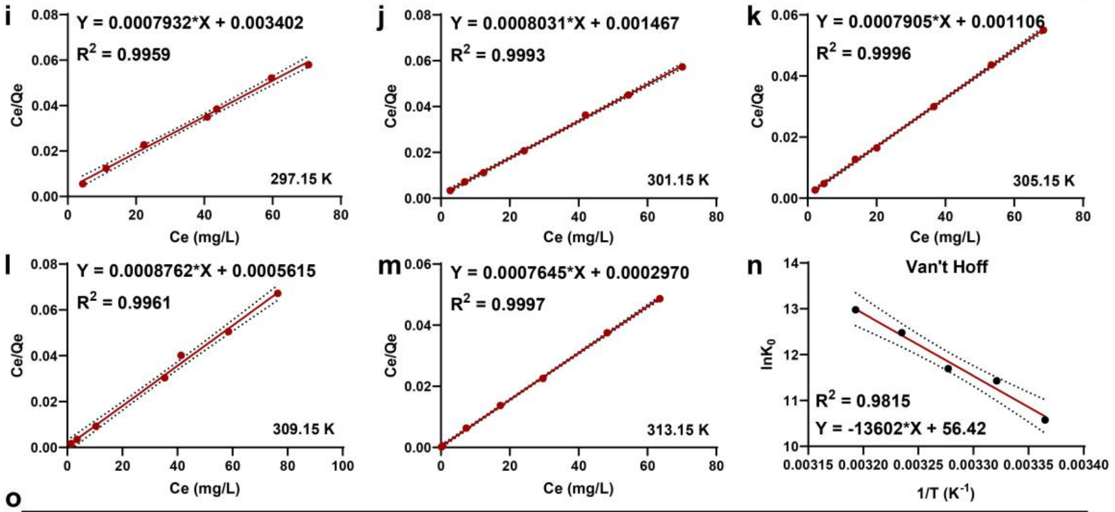


o

| T (K) | $lnK_0$ | $\Delta G^0$ (kJ $mol^{-1}$) | $\Delta H^0$ (kJ $mol^{-1}$) | $\Delta S^0$ (kJ $mol^{-1}$ $K^{-1}$) |
|---|---|---|---|---|
| 297.15 | 10.58 | -26.129 | | |
| 301.15 | 11.43 | -28.618 | | |
| 305.15 | 11.70 | -29.674 | 113.087 | 0.469 |
| 309.15 | 12.48 | -32.070 | | |
| 313.15 | 12.98 | -33.788 | | |

**Figure S6. The kinetics, isotherm, and thermodynamic studies of UA adsorption onto δ-$MnO_2$. a-d**, Adsorption kinetics study of UA adsorbed onto δ-$MnO_2$. (**a**) Adsorption capacity versus contact time of UA adsorption onto δ-$MnO_2$ in SIF at 37 ℃ (n = 3 replicates). The UA adsorption on δ-$MnO_2$ was found to be rapid in the early stages and subsequently became sluggish and stationary with increasing adsorption duration, achieving adsorption equilibrium at 4 h. (**b**,**c**) The linear form of pseudo-first-order kinetic (**b**) and pseudo-second-order kinetic (**c**) for the adsorption kinetics of UA onto δ-$MnO_2$. (**d**) Calculated constants of pseudo-first and second-order adsorption kinetic models for the adsorption of UA onto δ-$MnO_2$. Pseudo-second-order kinetic model has a better correlation value ($R^2$ = 0.9932) than pseudo-first-order kinetic model ($R^2$ = 0.9179). The calculated equilibrium adsorption capacity ($Q_e$, 1035 mg $g^{-1}$) of the pseudo-second-order kinetic model aligned well with the experimental $Q_e$ (~980 mg $g^{-1}$). **e-h**, The isotherm study of UA adsorbed onto δ-$MnO_2$. (**e**) Adsorption equilibrium ($Q_e$) versus equilibrium concentration ($C_e$) for the adsorption of UA onto δ-$MnO_2$ in SIF at 37 °C (n = 3 replicates). (**f**,**g**) The linear form of the Langmuir isotherm (**f**) and Freundlich isotherm (**g**) plots for adsorption of UA onto δ-$MnO_2$. (**h**) Calculated constants of the adsorption isotherm of the Langmuir and Freundlich equations for the adsorption of UA onto δ-$MnO_2$ at 310.15 K. The Langmuir model is applicable to homogenous surface and saturated monolayer adsorption. $R_L$ value between 0 and 1 imply a favorable adsorption of UA on δ-$MnO_2$. The Freundlich isotherm simulates multilayer adsorption and heterogeneous surfaces. The n value is in the range of 1-10, indicating that adsorption is favorable under the experimental conditions. The Langmuir model provides a superior match ($R^2$ = 0.9986) than Freundlich model ($R^2$ = 0.9896). **i-m**, Plots and linear regression of Ce/Qe vs. Ce for calculation of thermodynamic parameters about the adsorption of UA onto δ-$MnO_2$ at different temperatures including 297.15 (**i**), 301.15 (**j**), 305.15 (**k**), 309.15 (**l**), and 313.15 (**m**) K. **n**, Van’t Hoff plot for the adsorption of UA onto δ-$MnO_2$. The dashed line is 95% confidence interval of the linear regression. **o**, Calculated thermodynamic parameters for the adsorption of UA onto δ-$MnO_2$ at different temperatures.

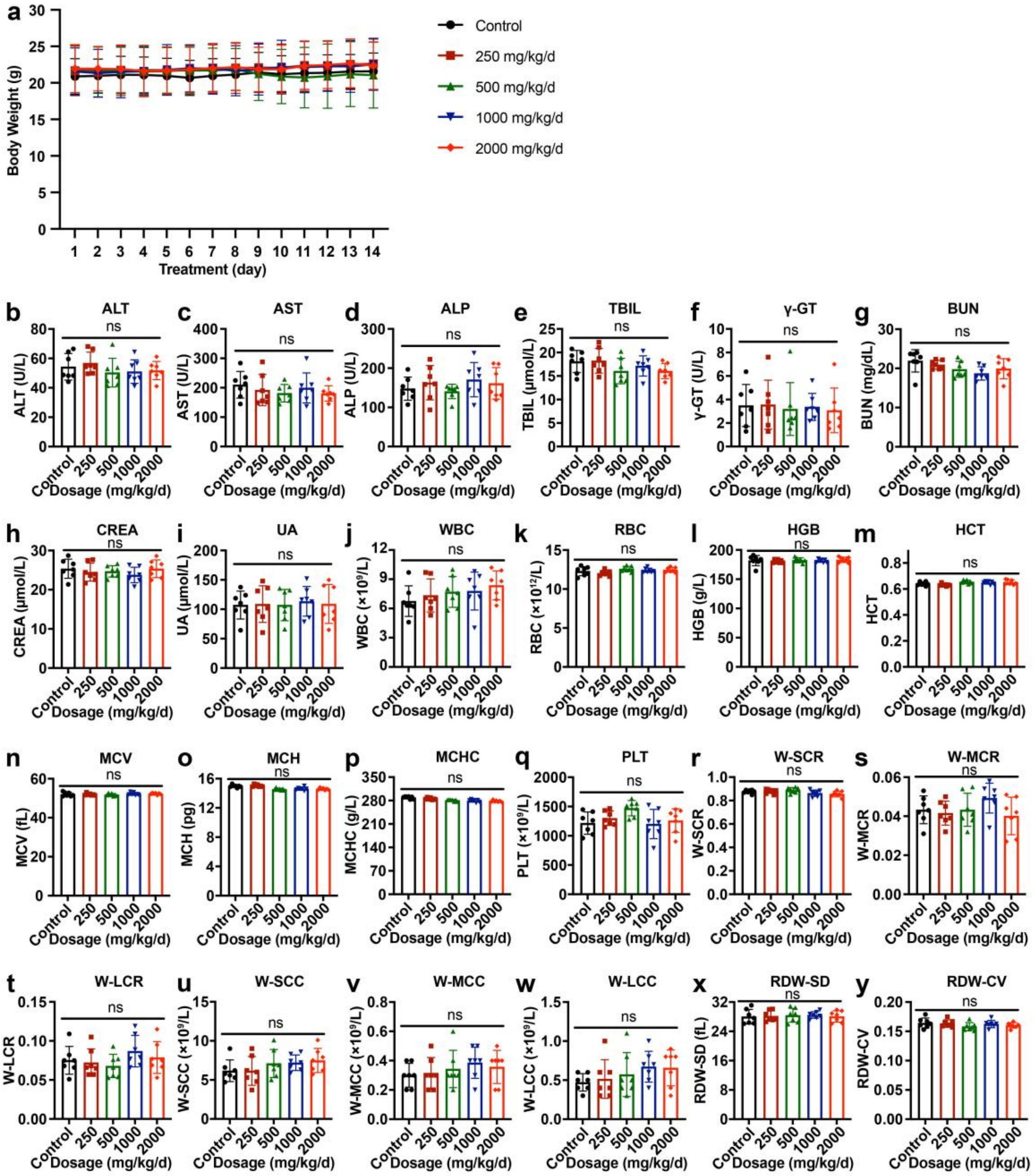


**Figure S7.** ***In vivo*** **safety assessment of δ-$MnO_2$. a**, The body weight changes of C57BL/6J mice after δ-$MnO_2$ oral administration for 14 days (n = 7). **b-i**, The analysis of serum biochemical indexes of liver (ALT, AST, ALP, TBIL, and γ-GT) and kidney (BUN, CREA, and UA) function after δ-$MnO_2$ oral administration for 14 days (n = 7). **j-y**, The routine blood examination of C57BL/6J mice after δ-$MnO_2$ oral administration for 14 days (n = 7).

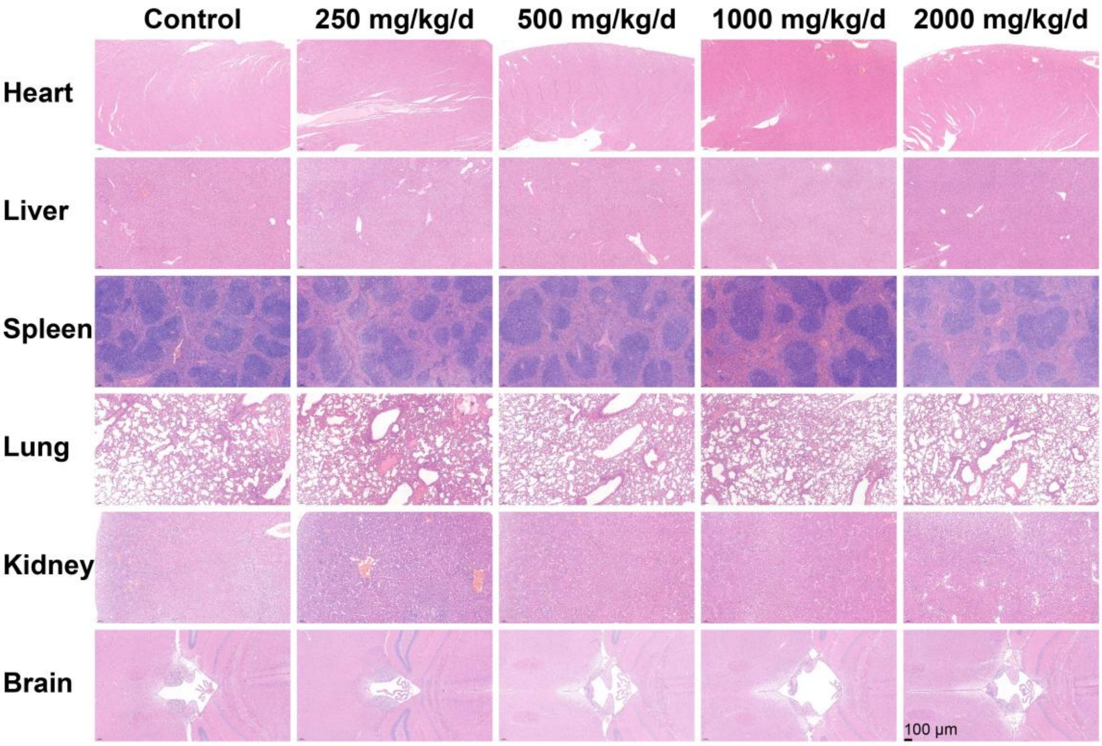


**Figure S8. The pathological analysis of major organs.** Representative hematoxylin and eosin (H&E) staining images of heart, liver, spleen, lung, kidney, and brain in mice treated with various dosages of δ-$MnO_2$ for 14 days. Scale bar: 100 μm.

**a**

| Independent variables | -α | Low level | Medium level | High level | +α |
|---|---|---|---|---|---|
| | -2 | -1 | 0 | +1 | +2 |
| Initial concentration (A, μM) | 200 | 300 | 400 | 500 | 600 |
| pH (B) | 5 | 6 | 7 | 8 | 9 |
| Adsorbent dose (C, g $L^{-1}$) | 0.05 | 0.06 | 0.07 | 0.08 | 0.09 |
| Contact time (D, min) | 15 | 30 | 45 | 60 | 75 |

**b**

| Std | Run | C (μM) | pH | A (g $L^{-1}$) | t (min) | Q (mg $g^{-1}$) |
|---|---|---|---|---|---|---|
| 26 | 1 | 400 | 7 | 0.07 | 45 | |
| 20 | 2 | 400 | 9 | 0.07 | 45 | |
| 17 | 3 | 200 | 7 | 0.07 | 45 | |
| 13 | 4 | 300 | 6 | 0.08 | 60 | |
| 28 | 5 | 400 | 7 | 0.07 | 45 | |
| 9 | 6 | 300 | 6 | 0.06 | 60 | |
| 30 | 7 | 400 | 7 | 0.07 | 45 | |
| 10 | 8 | 500 | 6 | 0.06 | 60 | |
| 1 | 9 | 300 | 6 | 0.06 | 30 | |
| 14 | 10 | 500 | 6 | 0.08 | 60 | |
| 15 | 11 | 300 | 8 | 0.08 | 60 | |
| 8 | 12 | 500 | 8 | 0.08 | 30 | |
| 16 | 13 | 500 | 8 | 0.08 | 60 | |
| 19 | 14 | 400 | 5 | 0.07 | 45 | |
| 5 | 15 | 300 | 6 | 0.08 | 30 | |
| 27 | 16 | 400 | 7 | 0.07 | 45 | |
| 6 | 17 | 500 | 6 | 0.08 | 30 | |
| 2 | 18 | 500 | 6 | 0.06 | 30 | |
| 25 | 19 | 400 | 7 | 0.07 | 45 | |
| 18 | 20 | 600 | 7 | 0.07 | 45 | |
| 22 | 21 | 400 | 7 | 0.09 | 45 | |
| 23 | 22 | 400 | 7 | 0.07 | 15 | |
| 11 | 23 | 300 | 8 | 0.06 | 60 | |
| 7 | 24 | 300 | 8 | 0.08 | 30 | |
| 24 | 25 | 400 | 7 | 0.07 | 75 | |
| 4 | 26 | 500 | 8 | 0.06 | 30 | |
| 29 | 27 | 400 | 7 | 0.07 | 45 | |
| 3 | 28 | 300 | 8 | 0.06 | 30 | |
| 12 | 29 | 500 | 8 | 0.06 | 60 | |
| 21 | 30 | 400 | 7 | 0.05 | 45 | |

**Figure S9. The parameters of RSM-CCD study. a**, The coded values of independent variables for the RSM-CCD factor levels in the experiment design for optimization of UA adsorption condition. **b**, The actual values in the experimental design data matrix for optimization of UA adsorption condition using RSM-CCD.
